# Metabolic modeling of microbial communities in the chicken ceca reveals a landscape of competition and co-operation

**DOI:** 10.1101/2024.10.14.618310

**Authors:** Irina Utkina, Yi Fan, Benjamin P. Willing, John Parkinson

**Affiliations:** Department of Molecular Genetics, University of Toronto, Toronto ON, Canada; Program in Molecular Medicine, Hospital for Sick Children, Toronto ON, Canada; Department of Agricultural, Food and Nutritional Science, University of Alberta, Edmonton, Canada; Department of Biochemistry, University of Toronto, Toronto ON, Canada

**Keywords:** metabolic modeling, keystone species, community assembly, poultry microbiome, cross-feeding

## Abstract

With their ability to degrade dietary fibers to liberate otherwise unavailable substrates, members of the Bacteroidales exert a substantial influence on the microbiome of the lower intestine. Currently our knowledge of how this influence translates to the metabolic interactions that support community structure is limited. Here we applied constraints-based modeling to chicken cecal communities to investigate metabolic interactions in the presence and absence of *Bacteroides*. From metagenomic datasets previously generated from 33 chicken ceca, we constructed 237 metagenome-assembled genomes. Metabolic modeling of communities built from these genomes generated profiles of short chain fatty acids largely consistent with experimental assays and confirmed the role of *B. fragilis* as a metabolic hub, central to the production of metabolites consumed by other taxa. In its absence, communities undergo significant functional reconfiguration, with metabolic roles typically fulfilled by *B. fragilis* assumed by multiple taxa. Beyond *B. fragilis*, we found *Escherichia coli* and *Lactobacillus crispatus* also mediate influential metabolic roles that vary in the presence or absence of *B. fragilis*. Compensatory adaptations adopted by the microbiome in the absence of *B. fragilis* resulted in metabolic profiles previously associated with inflammatory bowel disease in humans, including energy deficiency, increased lactate production and altered amino acid metabolism. This work demonstrates the potential of chicken cecal microbiomes to investigate the complex metabolic interactions and key contributions that drive community dynamics.

## INTRODUCTION

The gut microbiome consists of a complex, yet stable, community of microbes that confer beneficial roles to their host including the extraction of key nutrients, promoting gut health and limiting colonization by pathogens. Helping organize these communities are metabolic interactions involving the sharing (cross-feeding) of metabolites between taxa^1^. Many of these interactions are thought to be mediated by so-called keystone taxa which play a crucial role in shaping microbial community structure and function^2–7^. Within gut communities, keystone taxa are represented by genera such as *Bacteroides* and *Lactobacillus* that have been shown to exert considerable influence within their communities^8,9^. For example, while *Bacteroides* spp. play an important role in the degradation of complex carbohydrates and production of short chain fatty acids (SCFAs)^10,11^, *Lactobacillus* spp. have been identified as divisive members, characterized by both positive and negative interactions with other microbes^12^. Currently our understanding of the cross-feeding relationships that drive community dynamics and the contribution of keystone taxa is limited.

Community metabolic modeling offers a powerful approach to unravel the complex interactions within the gut microbiome. Integration of genome-scale metabolic models from multiple microbes within a community-scale framework provides a mechanistic basis to dissect the metabolic landscape of microbial communities and the contributions of individual taxa^13–15^. Recent advances in community-scale metabolic modeling have enabled predictions of the production of key metabolites such as SCFAs, which help support gut health^16,17^ as well as identification of potential cross-feeding interactions between bacterial species^13–15^ and changes in the concentrations of metabolites associated with a variety of human diseases^18,19^. Beyond human health, the gut microbiome has attracted considerable interest from the livestock industry. For example, with global bans on the use of antibiotic growth promoters (AGPs)^20^ attention is turning to manipulating gut communities in chickens to address issues of food insecurity associated with a surge in enteric infections^20^.

In this study, we leverage community metabolic modeling to investigate the influence of keystone taxa on microbial communities associated with the chicken cecum, a major site of digestion of insoluble fibers and fermentation and host to the most diverse community in the poultry gut. Applying *in silico* modelling based on previously published metagenomic data, we investigate the impact of *Bacteroides fragilis* on community and metabolic dynamics. Our models reveal that *B. fragilis* significantly impacts microbial diversity and structure of metabolic interactions. In *Bacteroides*-depleted communities, we observe an increase in the influence of other taxa including *Escherichia coli*, *Anaerostipes butyraticus*, and *Lactobacillus crispatus,* leading to a functional reorganization of the microbiome. Our findings highlight the pivotal role of the unique metabolic capabilities of *B. fragilis* and the potential implications for host health and productivity in its absence.

## RESULTS

### Bacteroides species influence the microbiome composition and functional capacity

To examine the metabolic influence of keystone taxa within cecal microbial communities we divided 33 samples from a recent study of 7-day old chickens^11^ into two groups: one characterized by the presence of *Bacteroides* species (HB, n=15), specifically *B. fragilis*, and the other without *Bacteroides* species (NB, n=18)^11^. Metagenomic datasets previously generated from these samples were used to generate metagenome-assembled genomes (MAGs). To enhance MAG quality and consistency across samples, we conducted both individual assemblies for each sample and a co-assembly of all samples. This combined approach allowed us to capture species present in individual samples and improve assembly quality for shared taxa by leveraging the combined data. Additionally, we re-assembled common taxa by pooling the corresponding reads from multiple samples (see Methods). This process yielded 237 high- and medium-quality MAGs (defined as having ≥50% completeness and ≤10% contamination) representing 227 taxa. Each MAG represented at least 0.5% relative abundance in at least one of the 33 samples (**Figure 1A**). Communities were largely dominated by representatives of families such as Lachnospiraceae, UBA660, and taxa belonging to Oscillospirales order, with significant variations in their relative abundances across samples (**Figure S1**). Next, we generated genome-scale metabolic models for each MAG, refining models through gap-filling and manual curation to ensure their ability to support growth on a corn feed diet and incorporate missing carbohydrate-active enzymes and fermentation pathways (see Methods). Subsequently, models were combined to reflect the cecal samples from which they were derived resulting in communities represented by 5 to 33 taxa with an average of 1,096±98 unique enzymes (as defined by enzyme commission numbers; ECs; **Figure 1A**). Further, each community contained from 2,187 to 3,368 (mean 3,001±280) unique reactions encompassing from 143 to 598 (mean 495±83) unique pathways (defined in MetaCyc^21^), with the number of unique pathways being significantly higher in NB communities compared to HB (p=0.0045, Wilcoxon Rank-Sum test; **Figure 1B**).

**Figure 1.**
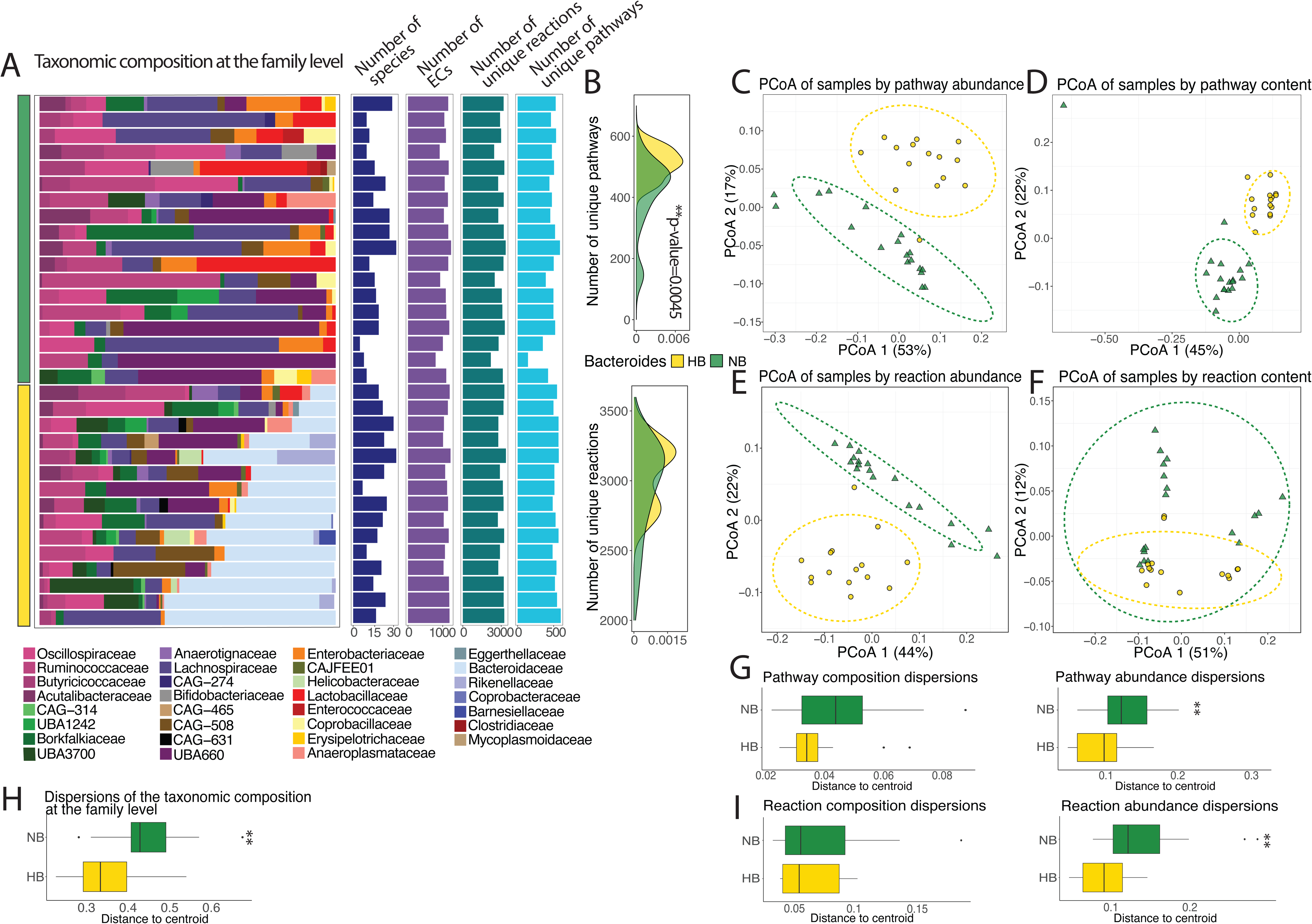
Taxonomic composition and functional analysis of chicken cecal microbial communities. **A.** Relative abundances of metagenome-assembled genomes (MAGs) consolidated by taxonomic family across 33 chicken cecal samples. The bar plots represent the numbers of unique species, ECs, unique reactions, and unique pathways based on metabolic model’s content. Samples are categorized into two groups: those with high Bacteroides abundance (HB, gold) and those without Bacteroides (NB, green). **B**. Distribution of numbers of unique metabolic reactions and pathways in NB and HB samples. Significance determined using Wilcoxon rank-sum test. **C, E**. PCoA of samples based on abundances of pathways (**C**) or reactions (**E**) content (i.e., reaction and pathway counts within each sample were summarized by weighing them by the relative abundance of the species contributing to that reaction or pathway), with dissimilarity matrices calculated using the Bray-Curtis index. **D, F**. PCoA of samples based on the presence or absence of pathways (**D**) or reactions (**F**) using Jaccard distances. **G-H.** Boxplots showing the distances from each stand to the centroid of the group that it belongs to. Distances were calculated from a PCoA of the Jaccard distances based on pathway (**G**, left) or reaction (**I**, left) content or PCoA of the Bray-Curtis dissimilarities based on pathway (**G**, right) or reaction (**I**, right) abundance; or relative abundances at the taxonomic family level (**H**). Significance determined using beta-dispersion (TukeyHSD) test. * - p-value < 0.05, ** - p-value <0.01, *** - p-value < 0.001.

Our reconstructed metabolic models revealed distinct metabolic profiles among taxa. *E. coli* had the largest number of enzymes, with broad representation across all pathways suggesting high functional diversity (**Figure S2; Table S1**), although we acknowledge this may also reflect its well annotated genome. *Bacteroides*, *Alistipes*, *Helicobacter*, *Barnesiella*, and *Coprobacter* spp., found exclusively in HB communities, exhibited expanded functionality in glycan and vitamin metabolism, suggesting they may play a dominant role in the breakdown of dietary fibers and vitamin synthesis. While most taxa possessed core functionalities across the major pathways, species within UBA1242 (e.g., *Caccovivens* spp.), UBA660 (e.g., *Faecenecus* and *Faecimonas* spp.), and CAG-508 (e.g., *Merdicola* spp.) families displayed sparser coverage of functions within carbohydrate metabolism. These microbial groups have been recently identified in diverse animal and human gut environments and are characterized by limited metabolic capabilities^22–27^. Despite their distant phylogenetic relationships, they each exhibit reduced genome sizes, diminished genetic repertoires, and numerous auxotrophies^28,29^. Further, these taxa exhibited minimal overlap with other species in amino acid metabolism capabilities, with distinct sets of enzymes involved in amino acid metabolism and xenobiotic biodegradation, suggesting they occupy a unique niche (e.g. mitigating environmental stressors) within the gut microbiome (**Figure S2)**.

Interestingly, while the prevalence of *Lactobacillus crispatus, Anaerostipes butyraticus* and *E. coli* in NB samples suggests that these taxa may become more dominant in the absence of *B. fragilis*, members of UBA660 family, specifically, *Faecenecus gallistercoris* and *Faecimonas intestinavium*, exhibit relatively high abundance (15-68%) in samples where the former three taxa are absent or exhibit low abundance (**Figure S1**). This may indicate competitive relationships, where less metabolically versatile taxa exploit specific substrates or metabolic niches that are less accessible to other bacteria. Members of the Oscillospirales order exhibited enhanced functionality in the metabolism of cofactors and vitamins, as well as carbohydrate metabolism, aligning with their roles in fiber degradation and SCFA production (**Figure S2**). These diverse metabolic capacities and interdependent roles of bacteria in the chicken cecal microbiome illustrate the complex relationships driving microbial community structure.

Principal Coordinate Analysis revealed that the presence of *B. fragilis* significantly influences both the taxonomic composition and metabolic potential of microbial communities (**Figures 1C - F**). NB and HB samples segregate on the basis of reaction and pathway content, regardless of similarity metric used. The high percentage of variance explained by the first two principal coordinates (70% for Bray-Curtis dissimilarity based on pathway abundance and 67% for Jaccard distance) together with a significant difference in metabolic pathway profiles (PERMANOVA p=0.001) highlights the impact of *B. fragilis* on community metabolism. However, the differences in within-group variability are not significant (TukeyHSD test p=0.13, permutest p=0.11) between the HB and NB groups (**Figure S3C**). Moreover, we observed distinct pathway and reaction abundance profiles between HB and NB groups (PERMANOVA p=0.001 for both), with greater variability within the NB communities (TukeyHSD p=0.015 and p=0.006, permutest p=0.009 and p=0.004, for pathways and reactions, respectively; **Figure S3B-C**). Overall, these findings show that *B. fragilis* has a major influence on metabolic interactions within the gut microbiome.

### Community metabolic modeling accurately predicts the production of acetate and propionate

To further dissect the metabolic contributions of *B. fragilis* within their communities we constructed *in silico* models for each of the 33 samples using the BacArena framework^30^ in which the relative abundance of each taxon was defined from the metagenomic datasets. Our *in silico* model was designed to mimic the physiochemical conditions of the chicken cecum, including low oxygen, neutral pH, peristalsis and release of cecal contents^29–32^ (see Methods). To assess the accuracy of our *in silico* ceca models, simulations were performed for 16 hours (representing a cycle of feeding), and predicted SCFA concentrations were compared with those previously measured experimentally^11^. We found a significant correlation between the predicted and experimentally derived concentrations for both acetate (spearman rho=0.53, p-value=0.0015; **Figure 2A**) and propionate (spearman rho=0.52, p-value=0.0021; **Figure 2A**). Moreover, the simulations reflected the experimentally measured differences for the NB and HB communities, although we note the model over-predicted the production of propionate by *Bacteroides* species (**Figure 2B**, **Figure S4**). The model was less predictive of butyrate production, potentially as a result of the model missing the impact of host epithelial cells which are known to uptake microbially derived butyrate^31,32^. Further, the discrepancy in the prediction of butyrate flux arises from the presence of butyrate-producing organisms (e.g. *Butyricicoccus* and *Faecalibacterium* spp.) in samples where the measured concentration was nearly zero. Previous studies have demonstrated cross-feeding interactions between *Bifidobacterium* spp. and *Faecalibacterium* spp., with acetate produced by the former enhancing butyrate production in the latter^33^. In our simulations, acetate produced by *Bifidobacterium pullorum* was utilized by other taxa, including *Butyricicoccus* and *Faecalibacterium* spp.^34–36^, to generate butyrate (**Figure S4**). This highlights the importance of studying metabolic interdependencies to better understand SCFA production in the gut microbiome. Overall, our models were able to accurately capture general trends and known dynamics of SCFA production, correctly predicting interactions between key gut microbes.

**Figure 2.**
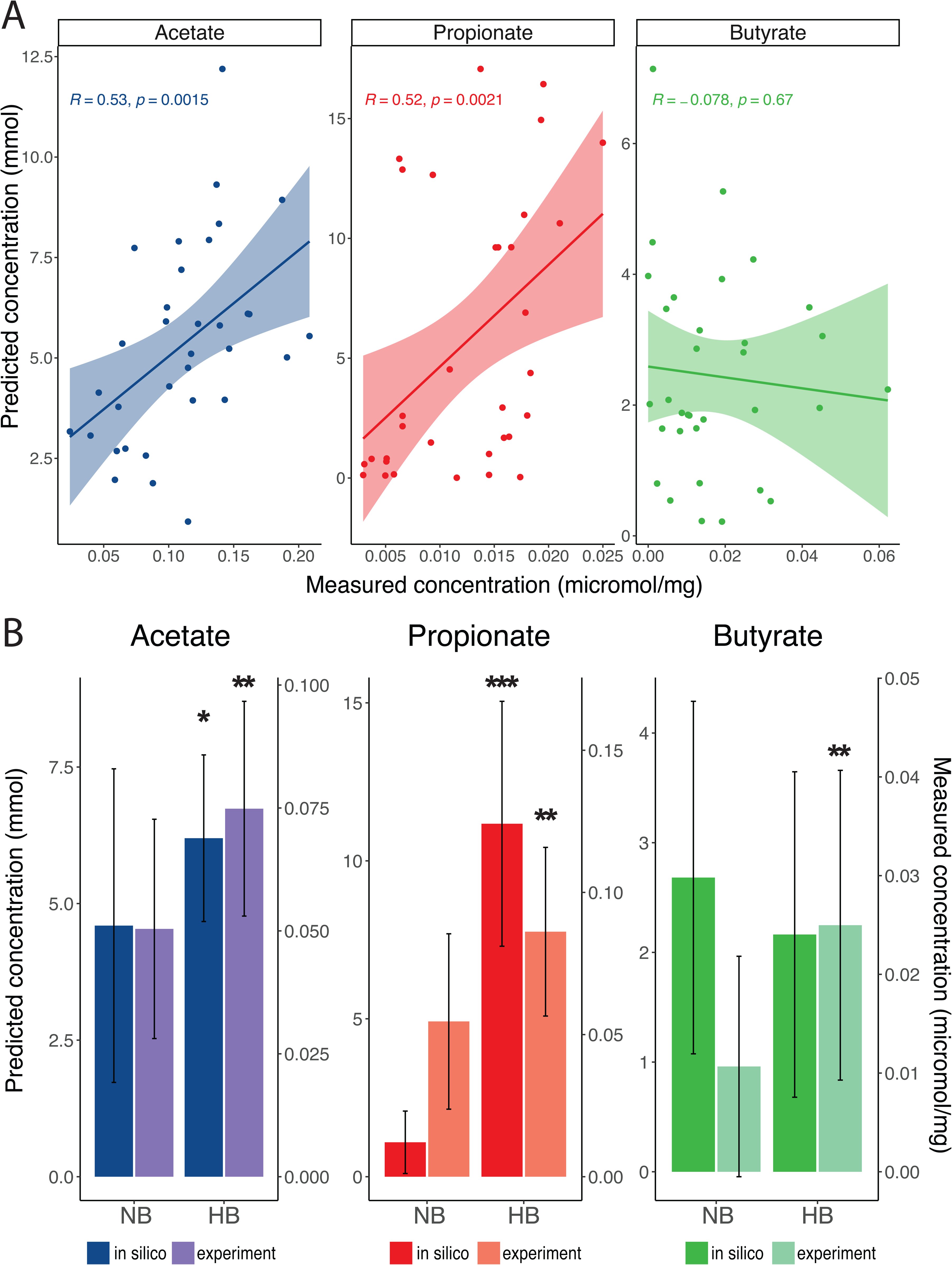
Comparison of predicted and measured SCFA concentrations in chicken cecal samples. **A**. Spearman rank correlation between *in silico* predicted and experimentally measured concentrations of acetate, propionate, and butyrate, with correlation coefficients (Spearman rho) and p-values indicated for each SCFA. Lines are fitted linear regressions, and colored areas around them indicate the 95% confidence interval. Each data point represents a sample. **B**. Bar plots comparing the average predicted and measured SCFA concentrations in samples with no Bacteroides (NB) and high (HB) Bacteroides abundance. The plots indicate the mean concentrations and standard errors for acetate, propionate, and butyrate with asterisks corresponding to the significance levels (Wilcoxon Rank-Sum test): * - p-value < 0.05, ** - p- value <0.01, *** - p-value < 0.001.

### *E. coli* and *B. fragilis* drive metabolic dynamics in the chicken cecal microbiome

Our analysis of metabolic interactions within chicken cecal microbial communities revealed distinct structural differences between HB and NB communities. To further investigate these differences, we used mutual information (MI) scores to construct a network based on metabolic overlap, where metabolic similarity refers to the extent of shared metabolic reactions between communities. We found that NB communities exhibited higher metabolic similarity than HB communities (pairwise MI scores 0.134±0.076 for NB and 0.123±0.068 for HB; **Figure 3A**). Although the difference in MI scores between NB and HB communities was not statistically significant (p=0.065, Wilcoxon Rank-Sum test), the trend suggests that the absence of *B*. *fragilis* leads to greater metabolic similarity. Since *Bacteroides* spp. play a fundamental role as primary fermenters and are often considered keystone taxa in their communities^37–40^, their potential monopoly of key resources may drive the reduction in community interactions and metabolic diversity. Moreover, the presence of both *B. fragilis* and *E. coli* resulted in further compartmentalization of metabolic functions, as evidenced by significantly lower overlap in metabolic functions (MI score 0.128±0.061) compared to NB communities with *E. coli* (MI score 0.154±0.08; p=0.022, Wilcoxon Rank-Sum test) and without *E. coli* (MI score 0.21±0.11; p=1e-05, Wilcoxon Rank-Sum test) (**Figure 3A**). Interestingly, samples lacking *E. coli* species exhibited highest pairwise MI scores (0.213±0.111 and 0.21±0.11, for HB and NB communities respectively) indicating that absence of *E. coli* promotes higher metabolic similarity irrespective of the presence or absence of *B. fragilis.* These findings suggest that consistent with its wide range of metabolic functions, *E. coli* acts as a metabolic generalist – a species capable of performing a broad range of metabolic activities, in contrast to specialists that have more limited, niche-specific roles **–** to fulfill roles vacated by other keystone taxa to support the metabolic versatility of the cecal microbiome (**Figure S2; Table S1**).

**Figure 3.**
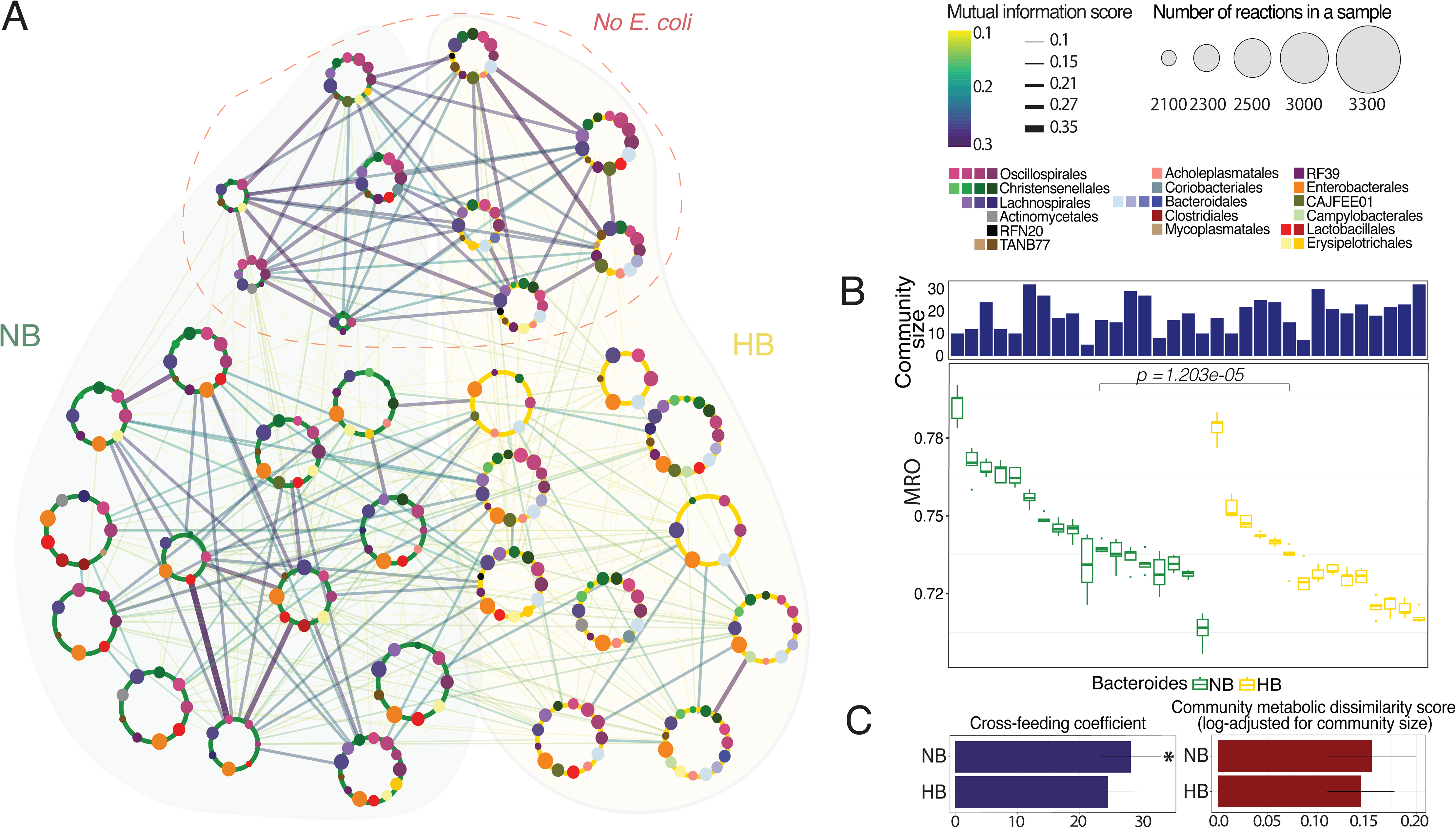
*E. coli* and *B. fragilis* influence metabolic similarity and cross-feeding in chicken cecal microbiomes. **A.** Mutual information-based network of chicken cecal microbial communities, highlighting metabolic similarities based on metabolic reaction content. Large nodes represent samples, with their sizes corresponding to the number of unique reactions in a sample, colored by category: no Bacteroides (NB, green) and high Bacteroides (HB, gold). Edges are colored and thickened according to the mutual information score, with thicker and darker edges indicating higher similarity. The top region outlined in red indicates samples lacking E. coli. Sizes of the smaller nodes within a sample, representing taxonomic families, correspond to the number of reactions covered by the metabolic models of species belonging to the respective taxonomic families (grouped by taxonomic order below). **B.** Bar plots showing community sizes (top) and box plots showing Metabolic Resource Overlap (MRO) within each sample (summarizing five *in silico* replicates). NB communities exhibit significantly higher MRO (p=1.203e-05, Wilcoxon Rank-Sum test) **C.** Cross-feeding coefficient (left) and Community Metabolic Dissimilarity (CMD) score (right) between NB and HB communities. NB communities show a significantly higher cross-feeding coefficient (*p=0.025) and a higher average CMD score.

While MI scores did not show a significant difference in metabolic similarity in NB and HB communities, the influence of additional taxa such as *E. coli* led us to investigate whether the absence of *B. fragilis* results in communities exhibiting a greater diversity of less dominant taxa, with increased competition and metabolic interactions. We applied three complementary metrics (see Methods): Metabolic Resource Overlap (MRO) score, a metric that quantifies both competition for shared resources as well as complementarity in metabolic capabilities^41^; Community Metabolic Dissimilarity (CMD) score, a metric that quantifies the diversity of metabolic functions in a community; and Cross-Feeding Coefficient (CFC) based on the number of metabolite exchanges. We found that NB communities had significantly higher MRO (p=1.2e- 05, Wilcoxon Rank-Sum test) and CFC (p=0.025, Wilcoxon Rank-Sum test) scores than HB communities (**Figure 3B-C**). While NB communities also featured higher CMD scores, consistent with reports showing that metabolic dissimilarities enhance cross-feeding between species^42^, this difference was not significant. Overall, these findings show that NB communities exhibit greater metabolic similarity between communities, along with enhanced cross-feeding relationships and higher metabolic diversity within individual communities, supporting the hypothesis that the presence of *B. fragilis* reduces metabolic diversity and interactions.

### Functional reorganization and emergence of alternate hubs in response to the absence of keystone taxa in the gut

Building upon the finding that the presence or absence of *B. fragilis* influences cross-feeding relationships among other taxa, we extended our analyses to examine changes in metabolite exchange dynamics in a larger set of taxa that could be considered metabolic hubs (defined here as taxa exchanging at least 12 metabolites with at least 10 other taxa across samples within each category; 87 taxa total - 40, 22 and 25 taxa associated with NB only, HB only and both, respectively). These taxa were used to generate profiles of interactions with other taxa and subsequently clustered to identify cliques of cross-feeding taxa (**Figure 4A**; **Table S2**). Consistent with our previous findings *B. fragilis* represents the central hub in the HB communities, while in its absence, *E. coli, A. butyraticus* and *L. crispatus,* adopt a greater range of metabolic interactions. The shift towards a more partitioned network observed in NB communities was further characterized by the formation of distinct cliques, representing groups of taxa engaging in mutual cross-feeding interactions. Within each clique, a single taxon (e.g. *Mediterraneibacter sp904377845, Lactobacillus johnsonii* and *Borkfalkia sp944368795*) appears to dominate interactions acting as a metabolic hub for each group of taxa.

**Figure 4.**
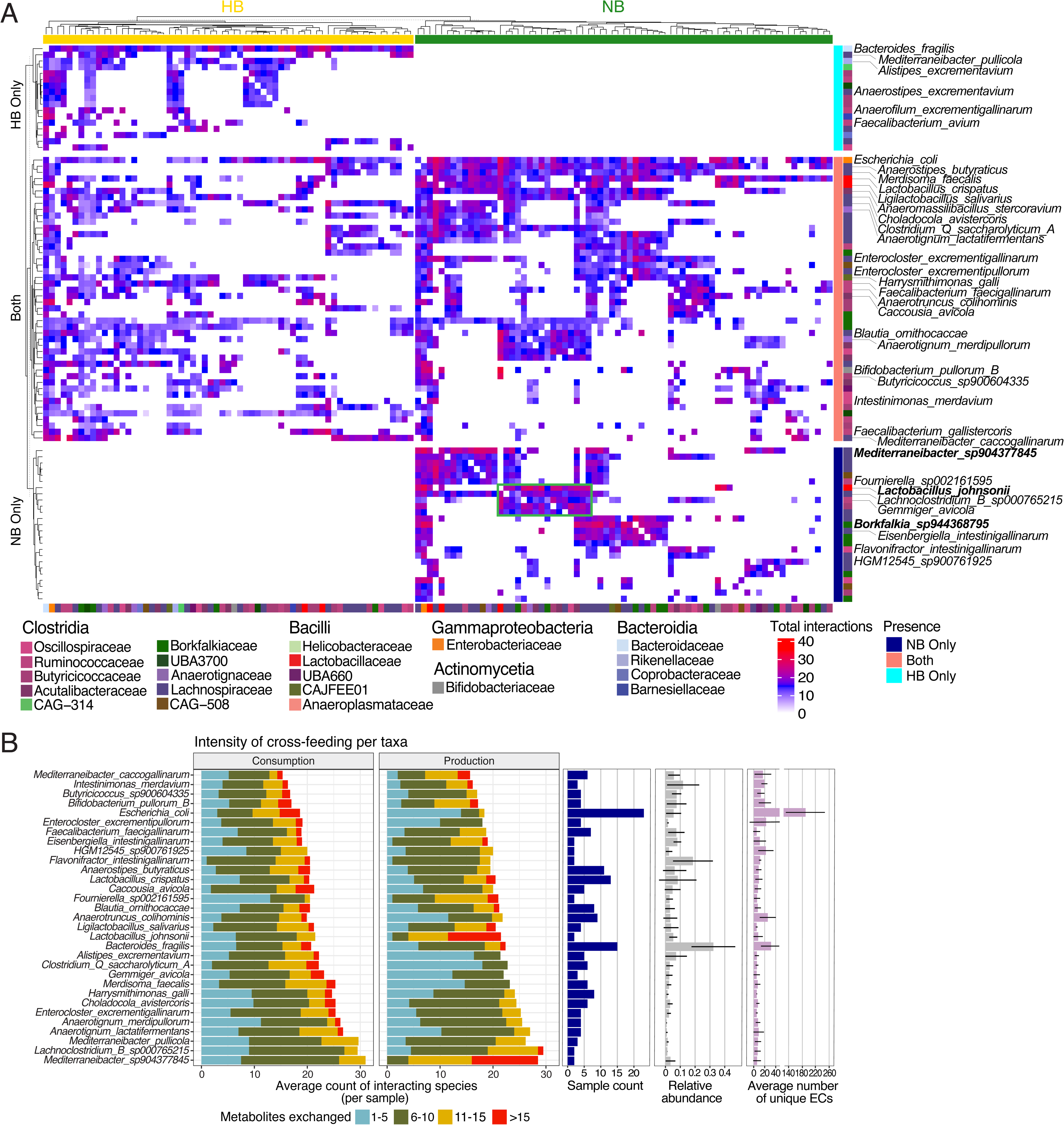
Major metabolic hubs and cross-feeding dynamics in HB and NB microbial communities. **A**. Heatmap of interactions between species (clustered hierarchically using complete linkage and euclidean distance, with rows split by HB/NB presence pattern and columns split by sample type (HB/NB), with clustering performed within each split), highlighting top cross-feeders defined as those forming at least 10 cross-feeding pairs with the exchange of a minimum of 12 metabolites across samples within each category, HB or NB. The interactions observed across multiple samples within each category were normalized by the number of samples in which each cross-feeding pair was present. Key taxa are labeled in this heatmap, including taxa present in at least two samples, forming an average of at least 230 interactions per sample and possessing on average at least 5 unique EC per sample. The tiles in the heatmap are colored based on the normalized total number of interactions for each species pair, with a color gradient indicating the interaction intensity. Blocks of taxon interactions represent ’cliques’, indicating groups of species with frequent metabolic exchanges. For example, the green-highlighted clique shows *Lactobacillus johnsonii* acting as a ’metabolic hub,’ engaging in extensive interactions with taxa from the families Lachnospiraceae (*Lachnoclostridium B sp000765215*, *Fusicatenibacter sp900543115*, *Blautia ornithocaccae*, *Eisenbergiella sp904392525*, *Choladocola avistercoris*, *Merdisoma faecalis*), Acutalibacteriaceae (*Eubacterium R faecavium*, *Acutalibacter ornithocaccae*, *Anaeromassilibacillus stercoravium*), Borkfalkiaceae (*Coproplasma stercoravium*, *Coproplasma stercorigallinarum*), Anaerotignaceae (*Anaerotignum merdipullorum*), Butyricicoccaceae (*Butyricicoccus avicola*), Ruminococcaceae (*Gemmiger avicola*), and Oscillospiraceae (*Dysosmobacter pullicola*), as well as with each other. Taxa mentioned in the main text are highlighted in bold. **B**. Stacked bar charts representing the intensity of cross-feeding per taxon. The bars indicate the average number of interacting species for each taxon per sample, divided into metabolite consumption (left) and production (right). Colors correspond to the number of metabolites exchanged, with categories ranging from 1-5, 6- 10, 11-15, and >15 metabolites. The bar plots to the right represent the sample count (number of samples in which each species is present, blue), relative abundance (grey), and the average number of unique enzymes (ECs) taxon possesses per sample (purple). Subset of taxa displayed is present in at least two samples, exhibit an average of at least 230 total interactions per sample, and possess a minimum of 5 unique EC numbers per sample.

To further clarify how ‘metabolic hub’ taxa exert their influence within their communities, we examined their interactions with other taxa in the context of prevalence across samples, relative abundance, and provision of unique functionality (**Figure 4B**, **Figure S5**). Surprisingly, a taxon’s unique metabolic potential within its community (as estimated by the number of unique enzymes) did not correlate with its number of metabolic interactions (**Figure S5A**) although we did observe a positive correlation with prevalence (**Figure S5B**). These findings suggest that while the number of unique functions a taxon brings to a community might increase its ability to engraft, the community does not exploit that additional functionality. Indeed, *E. coli*, which is both prevalent and possesses the greatest set of unique functions, appears to benefit in its community from the consumption of many metabolites from multiple species with only modest contributions to production (**Figure 4B**). Other prevalent taxa exhibiting a broad range of interactions, such as *L. crispatus* and *A. butyraticus,* with fewer unique enzymes did not exhibit the same disparity in consumption and production. At the other end of the spectrum, we found some less prevalent taxa, present in only a limited number of samples (e.g., *L. johnsonii* and *M. sp904377845*) exert their influence by specializing in the production of a wide array of metabolites utilized by a large number of other taxa (**Figure 4B**).

These findings reveal a hierarchy of taxa acting as metabolic hubs, where taxa such as *B. fragilis* and *E. coli* are responsible for driving the bulk of interactions, supplemented by additional hubs (e.g., *A. butyraticus* and *L. crispatus*). Further, within their communities, there exist cliques of taxa sharing metabolic interactions, organized around relatively rare taxa. This hierarchy highlights the varying abilities of each type of hub taxa to colonize within their communities, with *B. fragilis* and *E. coli* capable of colonizing many communities and establishing extensive cross-feeding networks. Others such as *M. sp904377845* and *L. johnsonii*, appear more limited in their ability to colonize communities and feature a smaller range of interactions with only a subset of taxa.

### Role of key metabolic hubs in modulating cross-feeding interactions

To further assess how the presence/absence of different metabolic hubs may impact cross-feeding relationships within their communities, we constructed a linear model based on the abundance of *E. coli* and the presence/absence of three additional metabolic hub taxa (*A. butyraticus, L. crispatus* and *B. fragilis*; see Methods). A relatively high abundance (>5%) of *E. coli* abundance was associated with increased cross-feeding in *Bacteroides*-depleted communities, potentially indicating the ability of *E. coli* to replace community functions otherwise performed by *B. fragilis* (**Figure S5C; Table S3**). Interestingly, only one of the 15 HB samples exhibited such an abundance, indicating competition between these two taxa. Extending this analysis to two other metabolic hub taxa (*A. butyraticus* and *L. crispatus*), we found their presence in the absence of *E. coli* was associated with increased cross-feeding activity, suggesting their ability to provide an alternative source of metabolic functions (**Figure S5D-E; Table S3**). Communities featuring both *E. coli* together with *A.butyraticus* displayed lower cross-feeding activity, reflecting potential competition between these taxa. In contrast, NB communities with *L. crispatus* showed higher cross-feeding activity compared to HB communities, regardless of *E. coli* abundance, indicating an antagonistic relationship between *L. crispatus* and *Bacteroides*.

Since our linear model focused only on taxa representing metabolic hubs, we were interested in how other metabolic hub taxa, specifically those mediating interactions within a subset of taxa (cliques), contribute to their communities. To avoid the confounding influence of dietary metabolites, we further restricted these analyses to metabolites only produced by bacteria. As before, *B. fragilis* represents the central hub in the HB communities, while *E. coli, A. butyraticus and L. crispatus* assume additional interactions in NB communities (**Figure S6A; Table S4**). Moreover, we also observed the formation of metabolic cliques, with greater numbers of interactions in NB communities. Comparing the eight largest cliques specific to either NB or HB communities (labelled A-H), we identified 48 unique microbially-derived metabolites exchanged within these cliques (ranging from 24 in clique H to 37 in clique A). While taxa associated with these cliques may not necessarily be present in the same samples, these cliques nonetheless represent groups of taxa that occupy a similar metabolic niche – that is, they share specific sets of metabolic functions and interactions they engage in, contributing to similar ecological roles across different communities.

This aligns with previous research demonstrating enrichment of mutualistic interactions in co-occurring microbial communities^41^. While there is a substantial overlap in metabolites exchanged across the eight cliques investigated – 17 metabolites were common to all cliques, and an additional 16 were shared among at least five cliques (**Figure S6B**) – individual cliques exhibit unique metabolic exchanges reflecting the specific taxa involved. This suggest that, despite a shared core of metabolic pathways and intermediates, each clique possesses distinct metabolic interactions driven by its constituent taxa. For instance, in the NB-specific clique B, *Bifidobacterium* spp. And members of Lactobacillaceae uniquely produced and exchanged D-alanine, melibiose, and spermidine (**Figure S6A-B**). Relative to other cliques, clique D, comprising members of Oscillospirales, Lachnospirales, Christensenellales, Coriobacteriales, and RF39, exhibited a more limited range of metabolite exchanges, lacking common fermentation products but uniquely exchanging NAD, highlighting the role of redox reactions and energy metabolism within this clique. These findings highlight how specific taxa contribute to functional diversity within microbial communities by occupying distinct metabolic niches, thereby shaping the overall metabolic landscape of their environments.

The changes in cross-feeding relationships associated with the absence of *B. fragilis* led us to investigate how this translates in terms of exchange of specific metabolites. We therefore analyzed the production and consumption of 18 metabolites selected on the basis of their relationship with 4 keystone taxa (see Methods; **Figure 5**). As expected, in HB communities *B. fragilis* is the predominant producer, contributing to the majority of the production of 10 metabolites, with *E. coli, Mediterraneibacter glycyrrhizinilyticus, Faecalibacterium gallistercoris,* and *Dysosmobacter pullicola* largely responsible for the production of a more limited set of metabolites including methanol, indole, uridine, formate and L-glutamine. In NB communities, the production of these metabolites is redistributed to multiple other species. For example, consistent with previous reports^33,36,43,44^, in the absence of *B. fragilis*, *E. coli* assumes the production of multiple metabolites otherwise produced by *B. fragilis* including acetate (along with members of *Clostridia*), ammonium (with *A. butyraticus*), succinate, propionate, sorbitol and GABA. These activities are supported through the production of lactate, uridine and L-glutamine by *L. crispatus*, the latter consistent with previous reports of its production^45^. In the presence of *B. fragilis*, *E. coli* alters its metabolic contributions to the generation of methanol, itself utilized by *B. fragilis.* Interestingly, in NB communities *A. butyraticus* adopts the role of methanol production (**Figure 5**).

**Figure 5.**
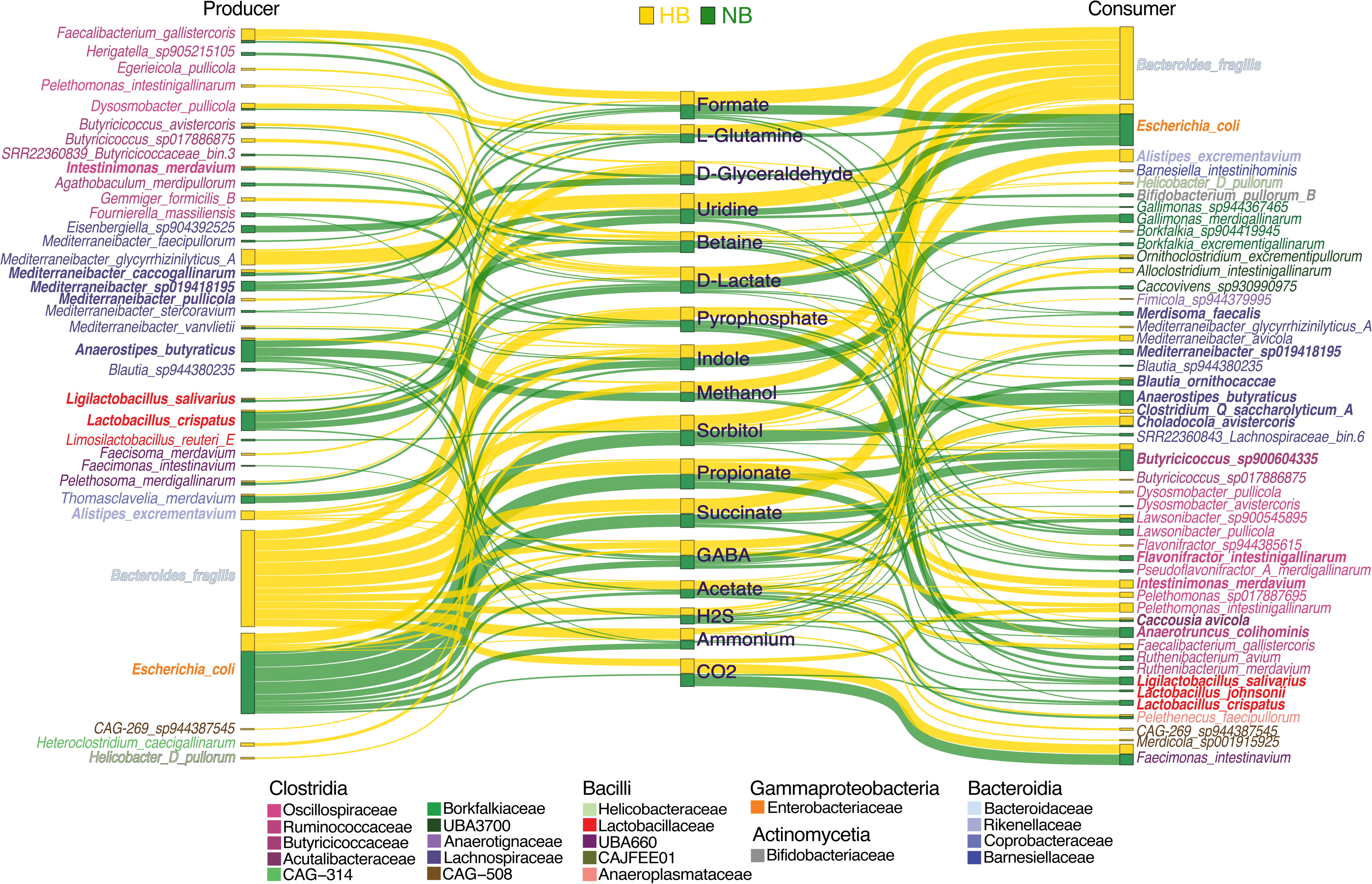
Shifts in cross-feeding networks: metabolic role redistribution in HB and NB communities. The Sankey diagram illustrates the cross-species interactions between primary producers and consumers of key cross-fed metabolites in the HB and NB communities. Producers are displayed on the left, cross-fed metabolites in the middle, and consumers on the right. Gold links represent contributions in HB communities, while green links represent those in NB communities. The thickness of each link is proportional to the contribution of the species (as a percentage) to the total production or consumption of each metabolite. To highlight the major players in cross-feeding dynamics, only the top three producers and consumers for each metabolite are shown, with a cutoff set at a minimum 5% contribution across samples within each category. Species highlighted in bold were previously described in Figure 4B. The diagram provides a visual summary of how key microbial taxa participate in the metabolic exchange, highlighting functional shifts between HB and NB communities.

Together our predictions show that in the absence of a keystone taxon (here *B. fragilis*), the cecal microbial community undergoes significant reorganization, characterized by the emergence of alternate metabolic hubs as well as a shift in metabolite production and consumption profiles by taxa present in both types of communities.

### Metabolic reorganization associated with the loss of *Bacteroides* in the chicken cecum reflects disease-associated metabolism in human IBD

Our comparisons of HB and NB communities, exhibit notable similarities to microbiome profiles associated with Inflammatory Bowel Disease (IBD) in humans (Ulcerative Colitis - UC and Crohn’s Disease - CD). Previous studies have reported shifts in the abundance and roles of both *B. fragilis* and *E. coli* – major metabolic hubs in our cecal communities – that correlate with disease states. The former being associated with reduced abundance in IBD patients, while the latter is associated with increased abundance^46–50^. Additionally, we observed increased abundance of *Alistipes* and *Faecalibacterium* species in HB samples (**Figure S1**), consistent with communities associated with healthy individuals in contrast to IBD patients^49,51,52^. Similarities between our study of HB and NB communities with healthy/IBD communities also extend to metabolic profiles. For example, the heterolactic fermentation pathway associated with lactic acid bacteria is more prevalent in NB communities, resulting in increased starch consumption and lactate production (**Figure 6**), consistent with elevated lactate levels observed in IBD patients^53–55^. NB communities are also associated with reduced production of propionate, a key source of energy for gut epithelial cells and a hallmark of IBD^55–58^. While studies have reported varied metabolic profiles in IBD, with both increased and decreased energy metabolism^55,59–61^, our findings align with the broader trend of altered energy metabolism in disease state.

**Figure 6.**
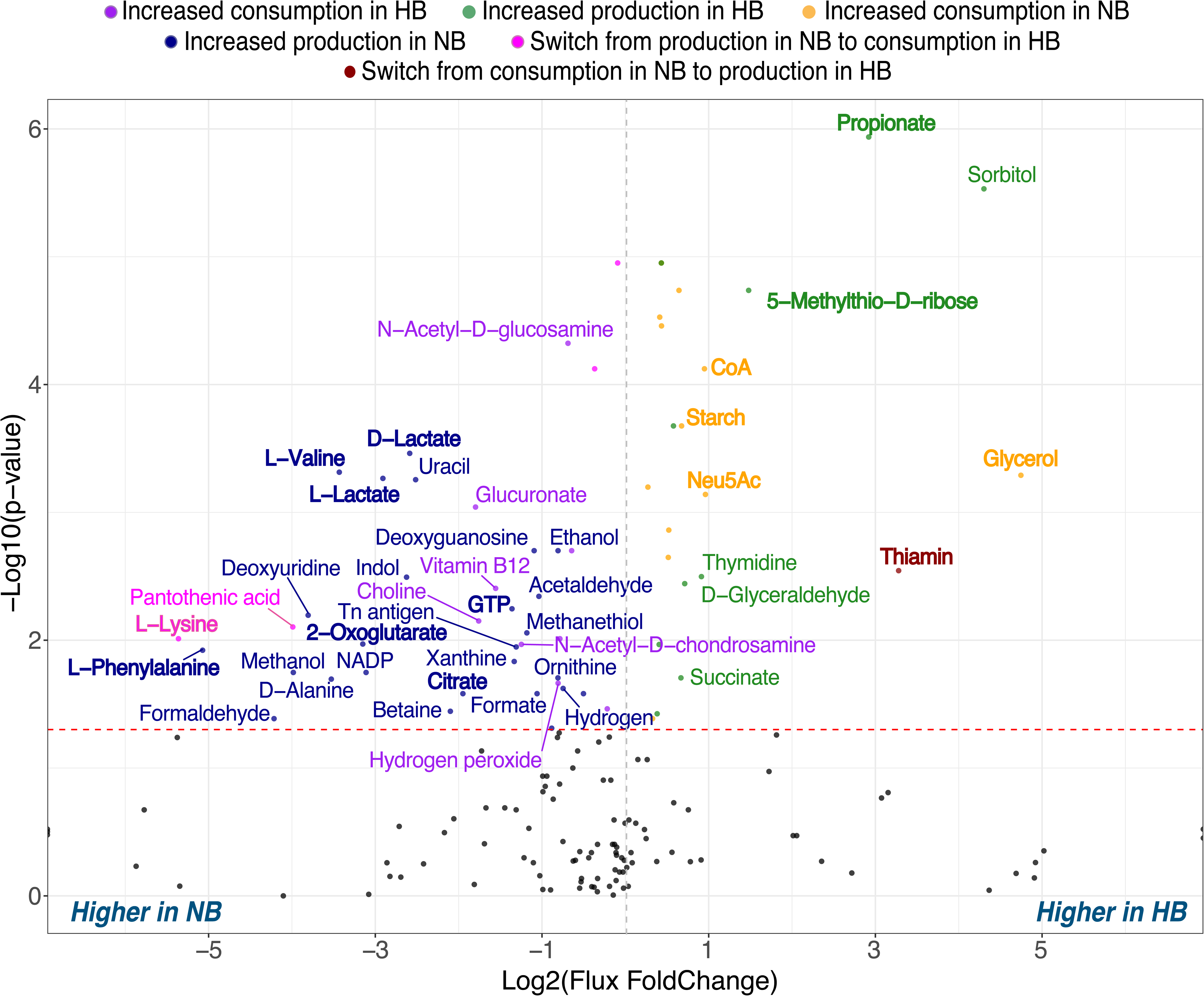
Comparison of metabolite exchange patterns between HB and NB communities. Each dot represents a metabolite being exchanged, with black dots indicating metabolites that do not show statistically significant changes in production or consumption between NB and HB groups. Statistically significant differences in total community fluxes (normalized to community size) were identified using the Wilcoxon rank-sum test. Dashed red line indicates the threshold for significance level, p-value < 0.05. Metabolites are color-coded based on the direction of their exchange: green - increased consumption in HB; blue - increased production in HB; orange - increased consumption in LB; purple - increased production in LB; red - switch from production in LB to consumption in HB; pink - switch from consumption in LB to production in HB. Metabolites with an absolute log2FoldChange greater than 0.7 are labeled and colored accordingly; metabolites discussed in results are highlighted in bold.

Specifically, we observed a metabolic shift in *Bacteroides*-depleted communities characterized by increased consumption of coenzyme A (CoA) and glycerol, alongside increased production of TCA cycle intermediates such as 2-oxoglutarate and citrate, as well as GTP. CoA, essential for fatty acid metabolism and the TCA cycle as an acyl group carrier, is being consumed more by the microbial community, suggesting upregulated lipid and energy metabolism pathways. However, the concurrent accumulation of TCA intermediates may indicate a bottleneck or dysregulation within the microbial TCA cycle, potentially due to limited CoA availability. This imbalance could reflect a shift towards fermentative metabolism in the absence of B. *fragilis*. Notably, reduced CoA levels have been observed in the colonic tissues of pig models and human patients with ulcerative colitis (UC), where they are associated with impaired fatty acid oxidation and energy metabolism in host cells^55,62^. While our findings pertain to increased microbial consumption of CoA, this could potentially decrease its availability in the gut lumen, impacting the host’s energy metabolism.

Beyond changes in energy metabolism, simulations also predict an increase in production of amino acids, including L-phenylalanine, L-valine and L-lysine in NB communities relative to HB (**Figure 6**), consistent with recent studies reporting alterations in levels of these amino acids in IBD patients^53,57,63,64^. Other notable metabolic similarities include reduced profiles of 5- Methylthio-D-ribose, a metabolite involved in the methionine salvage pathway, in NB communities, indicative of reduced sulfur cycle activity and consistent with deficiencies in sulfur-containing metabolites in microbial communities of IBD patients^18,19^. Thiamin, typically deficient in CD patients^53,65^, is predominantly produced by *B. fragilis*, and consequently reduced in NB communities. Finally, the increased consumption of sialic acid (Neu5Ac), a mucin-derived monosaccharide, by NB microbiomes suggests enhanced sialidase activity, which has been linked to intestinal inflammation and provides a growth advantage to bacteria like *E. coli*^66,67^. Thus, while the earlier study reported reduced inflammation in HB chicken cecal microbiomes^11^, the opposite may be inferred for NB microbiomes.

Overall, our findings reveal parallels between the community and metabolic dynamics of chicken cecal communities in the absence of *B. fragilis* and those associated with IBD, suggesting *Bacteroides-*rich communities may confer a protective role against inflammation.

## DISCUSSION

Previous studies suggest microbial communities in the gut display a hierarchical structure, with so-called keystone taxa acting as influential members through their ability to provide their communities with unique metabolic capacities^7,68–70^. The Bacteroidaceae family members are considered to be one of the most influential taxa, exerting a significant impact on metabolic interactions within their communities^37–39,71,72^. Previous studies have shown that the poultry gut, specifically the cecum, exhibits substantial fluctuations in the abundance of members of the Bacteroidaceae family, including the genus *Bacteroides*^11,73,74^. Notably, *Bacteroides* spp. are less abundant in intensively raised chickens compared to those raised in extensive systems (i.e., with access to the outdoors)^75^. Shifts in the abundance of *Bacteroides* can impact the community dynamics resulting in reduced feed efficiency, and heightened susceptibility to enteric diseases^76–78^. In humans, such shifts have also been associated with IBD^79,80^.

Here we leveraged a pre-existing dataset of metagenomic data from the cecal communities of chickens displaying a range of *Bacteroides* abundance and performed community metabolic modelling to dissect the influence of keystone taxa on metabolic interactions within their communities. Notably we found that our sample-specific community models accurately predicted the production SCFAs, particularly acetate and propionate, and recapitulated experimentally observed differences between *Bacteroides*-rich (HB) and -depleted communities (NB). Our findings demonstrate that the absence of *B. fragilis* results in more similar metabolic profiles across different microbiomes, while simultaneously increasing the range of metabolic functions within each individual community. However, this shift does not necessarily lead to an increase in overall microbial diversity. Moreover, NB communities exhibited significantly higher metabolic resource overlap within communities and increased cross-feeding interactions. This trend aligns with the role of *Bacteroides* spp. as keystone taxa, which typically dominate and monopolize key metabolic functions, thereby reducing metabolic diversity within their communities^81,82^.

By defining metabolic hubs as microbial taxa with a central role in metabolite exchange and distribution, we examined their influence on the overall metabolic landscape. While *B. fragilis* acts as a central hub, its absence triggers a shift towards a decentralized network characterized by distinct cross-feeding cliques and the emergence of new metabolic hubs, such as *E. coli*, *A. butyraticus*, and *L. crispatus*. Contrary to the expectation that a larger enzymatic repertoire directly translates to greater community influence, we observed a diverse range of strategies, from generalists like *E. coli,* which can perform a broad range of metabolic activities, to specialists like *L. johnsonii,* which focus on specific metabolic functions. Our analyses show that *E. coli* has a significant impact on cross-feeding, with increased activity when *E. coli* abundance exceeds 5% in *Bacteroides*-depleted communities. Interestingly, *L. johnsonii* and *L. crispatus,* previously described as eliciting a Marmite effect on the gut microbiome^12^, emerged as alternative metabolic hubs in the absence of *Bacteroides*, reflecting the antagonistic relationship between *Lactobacilli* and *Bacteroides* that has been observed in previous studies^11,83–85^. While *L. crispatus* had a positive effect on cross-feeding in the absence of *E. coli*, its presence in samples where *E. coli* was abundant led to a decrease in metabolic interactions. These observations highlight the shifting role of *L. crispatus* as it can both contribute to and limit the community’s metabolic interactions, depending on the presence of keystone taxa (e.g., *E. coli* and *B. fragilis*).

In our study we found that *B. fragilis* plays a central role in the production of metabolites utilized by other taxa whereas NB communities were characterized by a greater distribution of metabolic functions across a broader range of taxa. While this might be expected to contribute to community stability, similar changes in distributions are thought to contribute to disease associated with the human gut^86,87^. Indeed, our comparative analysis of HB and NB communities revealed striking parallels with IBD-affected human gut; NB communities exhibit increased lactate production, energy deficiency, and altered amino acid metabolism, alongside increased markers of inflammation. In contrast, HB communities displayed metabolic signatures associated with a healthy gut state including a higher production of SCFAs and essential vitamins, such as thiamin. Further, consistent with our cecal models, a decreased abundance of *Bacteroides*, particularly *B. fragilis*, which are typically abundant in healthy individuals, has been observed in patients with IBD, allowing opportunistic pathogens such as *E. coli* to flourish^46–50^. These findings highlight the potential of using models of chicken cecal communities to study gut microbial ecology and disease mechanisms.

This work establishes a framework for utilizing community metabolic modeling to study the poultry gut microbiome. Our approach facilitates the prediction of metabolic outputs, such as SCFA production, as well as elucidation of cross-feeding relationships. Moreover, our framework offers a mechanistic understanding of how keystone taxa, like *Bacteroides,* influence the ecological dynamics and metabolic landscape of the gut microbiome. Despite its promising results, we acknowledge our platform is not without some limitations. For example, community-level metabolic models may risk overemphasizing the metabolic roles of well characterized taxa, such as *E. coli*^88,89^. Further, the absence of host-microbiota interactions ignore the potential contribution of the host in terms of uptake of metabolites, as well as the influence of the host immune system. While the models accurately predict overall trends in metabolite production, they may overestimate or underestimate specific metabolic fluxes. This can be attributed to limitations in the functionality of some bacterial models, influenced in turn by the quality of the corresponding MAGs. Finally, the lack of data from upstream gut compartments may also limit our understanding of the precise metabolic environment in the cecum; here we assume that dietary components in amounts adjusted for known estimates of upstream absorption are directly available to members of the cecal community, ignoring the potential metabolic contributions from microbes present upstream of the ceca. Future research could integrate host factors, explore other gut regions, and assess the potential for targeted interventions to beneficially manipulate the microbiome.

## MATERIALS AND METHODS

### External data collection

The experimental methods and data collection procedures have been described previously^11^. Publicly available raw shotgun metagenomic sequence data from 33 chicken ceca and associated metadata associated with the study were downloaded from the sequence read archive (SRA; BioProject ID PRJNA902117).

### Chicken cecal metagenomic samples analysis

Metagenomic reads were subjected to quality filtering using BBTools/BBMap (v39.03)^90^. Single-sample assemblies and co-assemblies of short reads were conducted with MEGAHIT (v1.2.9)^91^, specifying a minimum contig length parameter of ‘–min-contig-len’ set to 1000. The Bowtie2 (v2.2.3)^92^ tool was employed to index assemblies and map filtered reads onto the contigs, creating abundance profiles. Following this, the metagenome assemblies underwent separate binning processes utilizing MetaBAT2 (v2.13)^93^ and CONCOCT (v1.1.0)^94^. Initially, samtools (v1.9)^95^ sorted the read mappings, and the read coverage was determined using the MetaBAT2 script ‘jgi_summarize_bam_contig_depths’. Subsequently, the binning process involved the ‘metabat2’ command with default settings for each assembly, using contig coverage across samples as input.

For CONCOCT, accessory scripts were employed to cut contigs into 10 kb fragments (cut_up_fasta.py), and the read coverage for these fragments was computed using CONCOCT_coverage_table.py. The fragment coverages were then utilized for binning the 10 kb fragments, and the clustered fragments underwent merging (merge_cutup_clustering.py) to generate the final CONCOCT bins (extra_fasta_bins.py). To dereplicate and consolidate the produced outputs from CONCOCT and MetaBAT2 binners, the metaWRAP (v1.2.3)^96^ ’bin refinement’ command was employed, utilizing parameters ’-x 10 -c 50’. These parameters signify a maximum bin contamination threshold of 10% and a minimum bin completeness threshold of 50%, as determined by CheckM^97^ estimates. Taxonomic annotation of generated bins was performed using GTDB-Tk (v2.3.2)^98^ command ‘gtdbtk classify_wf’ with default settings. Finally, the abundances of the draft genomes (bins) across the samples were quantified using the metaWRAP ‘quant_bins’ module.

To improve the consistency and quality of models representing the same species across different samples, different assembly strategies were employed depending on species prevalence and strain heterogeneity. Since all samples were derived from chickens raised in the same environment, we expected limited strain variation for most species, making it reasonable to pool reads from different samples. For species present in multiple samples (except E. coli), reads corresponding to each species were pooled from all samples where the specie was detected and reassembled, followed by re-binning using the previously described procedure to generate a single MAG per species (**Figure S3A;Table S5**). *E. coli* required a different approach due to its high prevalence (present in 23 samples) and strain heterogeneity. Initially, pooling all *E. coli* reads resulted in an 8.2Mb assembly, well above the typical *E. coli* genome size, suggesting the presence of multiple strains. To represent the multiple *E. coli* strains present across samples without retaining the 23 sample-specific *E. coli* individual assemblies with varying size and completeness, the bins derived from the individual assemblies were clustered based on average nucleotide identity using the fastANI software^99^. A distance matrix was constructed, and the bins were then clustered using the *hclust* function from the R-package stats. Five clusters were selected, and five cluster-specific *E. coli* strains were re-assembled, each representing a species pooled from samples within the corresponding cluster, followed by re-binning as described earlier. The resulting 5 MAGs vary in size from 4 to 5.8 Mb which is within the expected range for the *E. coli* genome^100^ (**Figure S3A**).

While most species benefited from sample-specific assembly followed by read pooling, five species yielded higher quality MAGs from the initial co-assembly of all 33 samples, based on completeness, contamination, and N50 metrics. These species were *Bacteroides fragilis*, *Anaerostipes butyraticus*, *Alistipes excrementavium*, *Limihabitans stercoravium*, and *Caccovivens* s*p944345705*. The characteristics and classification of the 237 reconstructed MAGs are provided in the **Table S5**.

### Model reconstruction and curation

The MAGs with a minimum relative abundance of 0.5% in each sample underwent reconstruction using gapseq v1.2 (commit 3e2d9393, Sequence DB md5sum: 17e92c9) with default settings^101^, resulting in a total of 233 metabolic models. Each sample was represented by 5 to 33 unique MAGs. Since the specific dietary composition used as chicken feed was not disclosed in the original study, the nutritional input for the models was formulated based on a typical chicken corn-based diet^102^ by translating molecular constituents into corresponding metabolites using vmh.org, foodb.ca and feedtables.com resources^103–105^. Due to limited information on fiber in the chicken diet, cellulose was utilized as a primary source of fiber according to available literature^106,107^. The absorption and digestion of metabolites in the upper gut compartments were taken into account when calculating the dietary input based on available information from studies on dietary absorption in poultry gut^108–113^. Additionally, urea and mucus were introduced to depict the cecal intestinal setting. Subsequently, this resulting nutritional composition served as an input medium for gapfilling step via gapseq. To ensure microbial degradation capabilities towards dietary components, profiling of carbohydrate-active enzymes (CAZymes) encoded within metagenomes used for model reconstruction was performed using dbCAN3^114^, which is based on the CAZy database - an expert-curated and routinely updated resource. Predicted CAZymes were filtered using an e-value threshold (>1e- 50) and required minimum 90% coverage with enzyme(s) assigned EC number(s). Predicted EC numbers along with substrates were mapped to respective reactions within modelSEED database and incorporated into models through gapseq adapt. Additionally, metabolic models of the species known as SCFA producers, were examined on the presence of fermentation pathways documented in the literature; missing pathways were added to the models.

### Quantitative PCR

qPCR of cecal digesta was performed using PercfeCTa SYBR green Supermix (Quantabio) with Primers used target the universal 16S rRNA gene: Forward CGGYCCAGACTCCTACGGG, Reverse TTACCGCGGCTGCTGGCAC^115^. The copy numbers were calculated back to copy numbers per gram of cecal digesta.

### Set up of *in silico* ceca and simulation of sample-specific community models

Microbial communities were simulated using the BacArena software^30^, which combines Flux Balance Analysis and individual-based modeling. BacArena represents each organism individually on a two-dimensional grid to model a spatial environment. Metabolites can be exchanged between individuals and diffuse in the environment, allowing for the dynamic update of the state of each microbe and its surroundings. The initial microbial community was seeded on a 100x100 cell grid. To mimic the flow of luminal contents in and out of the cecum while maintaining a relatively stable bacterial population, the following approach was implemented. Multiple studies have shown that the cecal contents are released only 2-3 times per day ^116,117^, which is a key feature for realistic simulation, considering the cecum’s role as a major hub for bacterial fermentation and SCFA production.

At each hourly iteration, metabolites representing the metabolic contents from the upstream compartment were added to the grid and evenly distributed between the grid cells, simulating environmental mixing and allowing access to dietary and bacterial-derived metabolites to all cells within the simulated environment. Every fourth iteration, a fraction of the cecum, including both metabolites and microbes, was randomly removed to mimic emptying of the cecum. The number of grid cells selected for removal of contents was determined to be 25% of the total to maintain realistic fluctuations in bacterial density and a generally consistent community size. To reflect physiological conditions of the chicken ceca, oxygen concentrations in simulations were set to 1e-08 mM to represent the nearly anaerobic environment of the chicken ceca, along with a pH of 7^118,119^.

The absolute abundances of microbial communities were measured via quantitative PCR (qPCR) assay, yielding copy numbers per gram of digesta ranging from 1.62e+09 to 3.02e+11. To reduce skewness and ensure smaller values were distinguishable while preventing larger values from dominating, abundance values were log-transformed. To preserve these differences in bacterial density between samples while making them computationally tractable, these values were then rescaled to range from 1000 to 5000 total individuals per community in each sample. This scaling approach maintains the proportional differences in bacterial density between samples, which is important for comparing metabolic outputs such as SCFA production across samples. At the same time, it ensures that even species with low abundances were represented by at least 5 individuals on the grid, while also allowing space for the community to grow and move within a grid of 10,000 cells. The communities were simulated in 5 replicates during 16 iterations, each representing an hour, to mimic the day of feeding.

### Comparison of functional capacities of community models

Sets of unique reactions and pathways within each sample-specific microbiome were extracted from the corresponding metabolic models comprising the sample. Binary matrices indicating the presence or absence of reactions and pathways in each sample were created for subsequent Jaccard distance calculations. Additionally, to quantify the metabolic potential within each sample while accounting for the varying contributions of different species, we summarized the reaction and pathway counts within each sample. Each reaction or pathway count was weighted by the relative abundance of the species contributing to that reaction or pathway. The resulting incidence matrices provided detailed information on the pathway and reaction abundance in the samples. Dissimilarity matrices were constructed from the incidence matrices described using *vegdist* function from the vegan R-package, and Principal Coordinate Analysis was performed using *cmdscale* function from stats R-package. Functional similarity networks detailing overlap between sample-specific microbiomes were constructed using mutual information score calculated via *mutinformation* function from infotheo R-package based on the metabolic reaction content of the samples. Networks were visualized using Cytoscape v.3.10.3^120^.

Community metabolic dissimilarity (CMD) score was calculated as the average of pairwise metabolic dissimilarities based on EC numbers within the community, adjusted for the community size. Specifically, the Bray-Curtis dissimilarity matrix for a sample was calculated from a presence/absence matrix where the columns represent species and the rows represent EC numbers, using the vegdist function from the vegan R-package. CMD score was then computed as a mean of the dissimilarity matrix, with further log-adjustment to account for the diminishing contribution of additional species in larger communities.

### Evaluation of cross-feeding and competition in *in silico* microbiomes

To evaluate the extent of metabolic competition within Bacteroides-rich and Bacteroides- depleted microbial communities, the metabolic resource overlap (MRO) between species was quantified as previously described ^41^. MRO is a metric that estimates the extent of overlap between the growth requirements of two or more metabolic models^41^. As such, it quantifies the potential for species to compete for a predetermined set of compounds in the media.

Cross-feeding interactions within microbial communities were investigated using metabolic reaction fluxes. We filtered negligible fluxes to retain absolute total flux values above 1e-06 millimoles per hour per gram of dry weight (mmol/gDW/hour) and categorized species as metabolite producers or consumers. By merging producer and consumer datasets on shared metabolites, cross-feeding interactions were summarized by counting and normalizing interaction occurrences across replicates within each category (based on *Bacteroides* presence). Key cross-feeders were identified using a threshold of at least 10 interactions involving the exchange of 12 or more distinct metabolites, focusing on species with substantial and consistent metabolic activity. Cross-feeding was quantified by retrieving all metabolite exchanges (import and export fluxes) during simulations and calculating the number of total interactions (cross-feeding pairs) in each sample. A cross-feeding coefficient was introduced as a metric of cross-feeding activity within a community, defined as the number of total cross-feeding pairs formed divided by all possible combinations of species pairs in the community. Interaction data were visualized using ComplexHeatmap R-package.

Next, the total production and consumption for each metabolite was calculated over the course of the simulation. The contribution of each species to the production and consumption was determined as a percentage of the total production/consumption. Exchanged metabolites were filtered to include only non-dietary metabolites, which are exclusively produced by bacteria. The pairwise interactions involving non-dietary metabolites summarized across samples are visualized as a heatmap in **Figure S6A** with raw data used to plot heatmap provided in **Table S3**.

To focus on metabolic niche adjustments involving keystone taxa, metabolites were further filtered based on the following criteria: 1) they are predominantly produced or consumed by keystone taxa, defined as those where at least 30% of produced metabolite is attributed to the production or consumption by a single keystone taxon; 2) to ensure the metabolite is involved in cross-feeding, at least 1% of the produced amount must be consumed by at least one other taxon; 3) metabolites produced in highest quantities by the same taxon in both HB and NB communities were excluded to emphasize shifts in metabolic roles. This filtering process resulted in the identification of 22 metabolites featuring different cross-feeding relationships in HB and NB comunities. Of these, 18 were selected for downstream visualization. Simple sugars D-Gluconate, D-Ribose, and D-Arabinose were excluded due to their relatively low rates of production; L-Lactate was excluded because its production and consumption patterns significantly overlapped with those of D-Lactate, so only D-Lactate is shown. . For each of these metabolites, the relative contributions of the top three producers and consumers were visualized (**Figure 5**). Sankey diagrams were generated using the networkD3 R-package. To highlight major contributors, a minimum contribution threshold of 5% was used to filter taxa for visualization.

To investigate differences in metabolite production and consumption between HB and NB microbial communities, the fluxes of exchanged metabolites were summarized within each community at every hour of the simulation and normalized by the total community size at that time point. Subsequently, the community fluxes (normalized by abundance) were aggregated over the 16-hour simulation period, and differences between the categories were evaluated using the Wilcoxon rank-sum statistical test. For graphical representation, metabolic fluxes were log2- transformed.

## Statistical analyses

All statistical analyses were performed in R v4.2.3. Significant differences for numbers of reactions and pathways, metabolite concentrations, metabolic fluxes, community metabolic diversity and cross-feeding activity coefficients were determined by non-parametric Mann– Whitney U-test (Wilcoxon rank-sum test). Spearman’s rank correlation coefficients and associated p-values were calculated between measured and predicted SCFA concentrations. Given *E. coli*’s high degree of connectivity, we explored its influence on metabolic exchanges within microbial communities. To evaluate how the presence or absence of this pivotal taxon determines the degree of cross-feeding activity, we constructed a linear model incorporating community size and the abundance of *E. coli* alongside key metabolic hubs: *A. butyraticus, L. crispatus,* and *B. fragilis* using the *lm* function in R. The response variable was the cross-feeding coefficient (CrossFeeding_coef), which quantifies the degree of metabolic interactions within the community. Predictor variables included categorical levels of *E. coli* abundance (Ecoli_category: "NoEcoli", "LowEcoli” (<5% relative abundance), "HighEcoli” (>5% relative abundance)), *Bacteroides* presence or absence that defines the community category (Bacteroides_presence: "NB" or "HB"), the presence of key taxa such as *Anaerostipes butyraticus* (Abutyraticus_presence) and *Lactobacillus crispatus* (Lcrispatus_presence), and community size (comm_size). Interaction terms between *E. coli* abundance and the other predictors were included to capture potential synergistic or antagonistic effects on cross-feeding. The model was specified as follows: *‘CrossFeeding_coef ∼ Ecoli_category:(Bacteroides_presence + Abutyraticus_presence + Lcrispatus_presence) + comm_size’.* The model summary is provided in the **Table S3**. All statistical analyses employed a significance threshold of p-value < 0.05.

## FUNDING SOURCES

This work was funded by grants from the Natural Sciences and Engineering Research Council of Canada (RGPIN-2019-06852) and the Ontario Research Fund-Research Excellence. High performance computing was provided by the SciNet HPC Consortium. SciNet is funded by Innovation, Science and Economic Development Canada; the Digital Research Alliance of Canada; the Ontario Research Fund: Research Excellence; and the University of Toronto. The funders had no role in study design, data collection and analysis, decision to publish, or preparation of the manuscript.

## AUTHOR CONTRIBUTIONS

J.P. and I.U. conceived and designed the study. I.U. performed simulations and data analysis. Y.F. performed qPCR experiments. B.P.W. provided critical reagents and contributed to data analysis. J.P. and I.U wrote the manuscript. All authors reviewed and/or edited the manuscript.

## COMPETING INTERESTS

The authors declare no competing interests.

## DATA AVAILABILITY STATEMENT

Microbial metabolic models and the source code used for simulations and data analysis have been deposited on GitHub (github.com/ParkinsonLab/Chicken_microbiome).

## SUPPLEMENTAL MATERIAL

**Supplementary Figure 1. Relative abundances of the common taxa across samples.** This heatmap illustrates the relative abundances of the top 52 most common taxa across 33 cecal samples, categorized into high *Bacteroides* (HB, in gold) and no *Bacteroides* (NB, in green) groups. Taxa present in at least four samples are shown, with the relative abundance represented by the color intensity of each tile.

**Supplementary Figure 2. Distribution of enzyme commission numbers (ECs) across metabolic models.** The columns of the heatmap represent individual ECs, while the rows correspond to metabolic models representing different taxa. ECs are annotated using KEGG pathway designations, indicated in two layers to reflect primary and secondary pathway associations. For ECs lacking KEGG pathway annotation, MetaCyc pathway annotation was utilized when available. For ECs mapped to 3 or more major pathway categories, “Multiple superpathways” category was assigned. Each row represents a different taxa, grouped by their presence in HB only, NB only, or both; color-coded.

**Supplementary Figure 3. Characteristics of 237 reconstructed MAGs. A.** Major characteristics of 237 MAGs reconstructed in this study: completeness, contamination, N50 and genome size. **B-D.** Boxplots showing the distances from each stand to the centroid of the group that it belongs to. Distances were calculated from a PCoA of the Jaccard distances based on reaction (**B**, left) or patway (**C**, left) content or PCoA of the Bray-Curtis dissimilarities based on reaction (**B**, right) or pathway (**C**, right) abundance; or relative abundances at the taxonomic family level (**D**). Significance determined using beta-dispersion (TukeyHSD) test. * - p-value < 0.05, ** - p-value <0.01, *** - p-value < 0.001.

**Supplementary Figure 4. SCFA reaction fluxes at the species-level.** Cumulative fluxes over 16 hours of simulation for acetate, propionate and butyrate, with samples (x-axis) ordered by the corresponding experimentally measured concentrations – from smallest to largest (from left to right). Colors correspond to the specie contributing to the production or consumption. Positive fluxes correspond to production, while negative fluxes indicated consumption.

**Supplementary Figure 5. Influence of species-specific metabolic potential and key taxa presence on cross-feeding activity. A-B.** Correlation plots showing the relationship between the average number of unique EC numbers presented in a specie within a sample and **(A)** the total number of interactions formed by a specie normalized by the number of samples where a specie is present (Pearson) **(B)** the number of samples where the specie is present (Pearson). Red line is fitted linear regression, and gray area indicates the 95% confidence interval. Each data point represents a species. Only species with at least 10 cross-feeding pairs involving at least 12 metabolites are shown. Selected species (present in at least two samples, above the threshold of 230 total interactions per sample and average number of unique EC per sample > 5) have labels that are color-coded corresponding to the taxonomic family they belong to. **C.** Effect of *E. coli* presence across both community categories (HB - High Bacteroides; NB - no Bacteroides) on cross-feeding activity, measured by the cross-feeding coefficient (CFC). Each dot corresponds to a CFC calculated for a single sample. Points and lines are color-coded based on *E. coli* abundance: red represents communities with high *E. coli* abundance (>5%), blue represents communities with low *E. coli* abundance (<5%), and gray represents communities where *E. coli* is absent. In the NB group, higher *E. coli* abundance is associated with a notable increase in cross-feeding activity compared to low or absent *E. coli*, suggesting a significant role for *E. coli* in enhancing cross-feeding, particularly in the absence of *Bacteroides*. Conversely, in HB communities, *E. coli* presence appears to have a more muted effect on cross-feeding, especially in low abundance. **D-E.** Interaction effects of *E. coli* and *A. butyraticus* **(D)** or *L. crispatus* **(E)** on cross-feeding activity, measured by the CFC, across categories. Different colors represent distinct interaction scenarios: *E. coli* & *A. butyraticus* **(D)/***L. crispatus* **(E)** absent (gray), *E. coli* absent & *A. butyraticus* **(D)/***L. crispatus* **(E)** present (cyan), low-abundance *E. coli* & *A. butyraticus* **(D)/***L. crispatus* **(E)** absent (blue), low-abundance *E. coli* & *A. butyraticus* **(D)/***L. crispatus* **(E)** present (orange), high-abundance *E. coli* & *A. butyraticus* **(D)/***L. crispatus* **(E)** absent (red), and high-abundance *E. coli* & *A. butyraticus* **(D)/***L. crispatus* **(E)** present (purple). Each point represents an average CFC for samples within each interaction scenario. NB communities with high *E. coli* abundance and without *A. butyraticus* (solid red line, **D**) show the highest cross-feeding activity. When *A. butyraticus* is present (dotted lines, **D**), the interaction patterns change: the cross-feeding coefficient decreases slightly in communities with high *E. coli*, but it increases in communities with low or no *E. coli*. This suggests that *A. butyraticus* may partly compensate for the absence or lower abundance of *E. coli*, contributing to cross-feeding, particularly in NB communities. The interaction between *A. butyraticus* and *E. coli* seems to reveal a potentially competitive relationship where the presence of both may reduce cross-feeding, while the presence of either one alone promotes cross-feeding in different ways. Presence of *L. crispatus* is associated with increased cross-feeding activity in NB compared to HB communities (**E**), regardless of *E. coli* abundance, while absence of *L. crispatus* doesn’t increase significantly the cross-feeding activity in NB when *E. coli* levels are low. This points to an impact of antagonistic relationship between *L. crispatus* and *Bacteroides* rather than potential competitive relationship between *L. crispatus* and *E. coli*.

**Supplementary Figure 6. Non-dietary metabolite exchanges across all 227 taxa in HB and NB communities. A.** The heatmap displays the normalized total interactions involving non-dietary metabolites exchanged between 227 taxa across 33 chicken cecum samples. Interactions were normalized by the number of samples in which each cross-feeding pair was present and are color-coded by interaction frequency. Taxa are color-coded based on their taxonomic families, interaction frequencies are indicated by color intensity. Taxa on y-axis are grouped by their presence in HB only, NB only, or both; color-coded. Distinct cliques of interacting species are highlighted in red boxes (A-H). **B.** The upset plot summarizes the unique and shared non-dietary metabolites exchanged within the investigated cliques (A-H) highlighted in panel A. Each bar represents the number of metabolites exchanged, with intersections indicating metabolites shared between multiple cliques. Some metabolite names have been shortened or modified for clarity in the figure. The following abbreviations are used: 2-Oxoglut for 2-Oxoglutarate, Ddca for dodecanoic acid, L-Asp for L-Aspartate, Glu for Glutamate, D-Lact and L-Lact for D- and L-Lactate, respectively, NAD for nicotinamide adenosine dinucleotide, Propion for propionate, Pyro-P for pyrophosphate, and Putrsc for putrescine.

**Supplementary Table 1. EC content in reconstructed metagenome assembled genomes annotated with KEGG and MetaCyc (related to Supplementary Figure 2)**

**Supplementary Table 2. Normalized cross-feeding interactions between top cross-feeding taxa in HB and NB communities (related to Figure 4A)**

**Supplementary Table 3. Linear model summary: impact of key metabolic hubs presence on cross-feeding activity.** The response variable is the cross-feeding coefficient, which quantifies the degree of metabolic interactions within the community. Predictor variables included categorical levels of *E. coli* abundance (Ecoli_category: "NoEcoli", "LowEcoli” (<5% relative abundance), "HighEcoli (>5% relative abundance)"), Bacteroides presence or absence that defines the community category (Bacteroides: "NB" or "HB"), the presence of key taxa such as Anaerostipes butyraticus (Abutyraticus_presence) and Lactobacillus crispatus (Lcrispatus_presence), and community size (comm_size).

**Supplementary Table 4. Non-dietary metabolite exchanges between taxa across HB and NB communities (related to Supplementary Figure 6).**

**Supplementary Table 5. Detailed characteristics and classification of the 237 reconstructed MAGs.**

## Notes

### Competing Interest Statement

The authors have declared no competing interest.

### Summary of Updates

Minor typos and mistakes revised; Methods section on Chicken cecal metagenomic samples analysis updated to clarify the process of metagenome-assemled genomes selection; Figure legends for Figure 1 and Figure 4 updated.

