## Supplementary figures and images for "Metabolic modeling of microbial communities in the chicken ceca reveals a landscape of competition and co-operation"

### Supplemental Figure 1

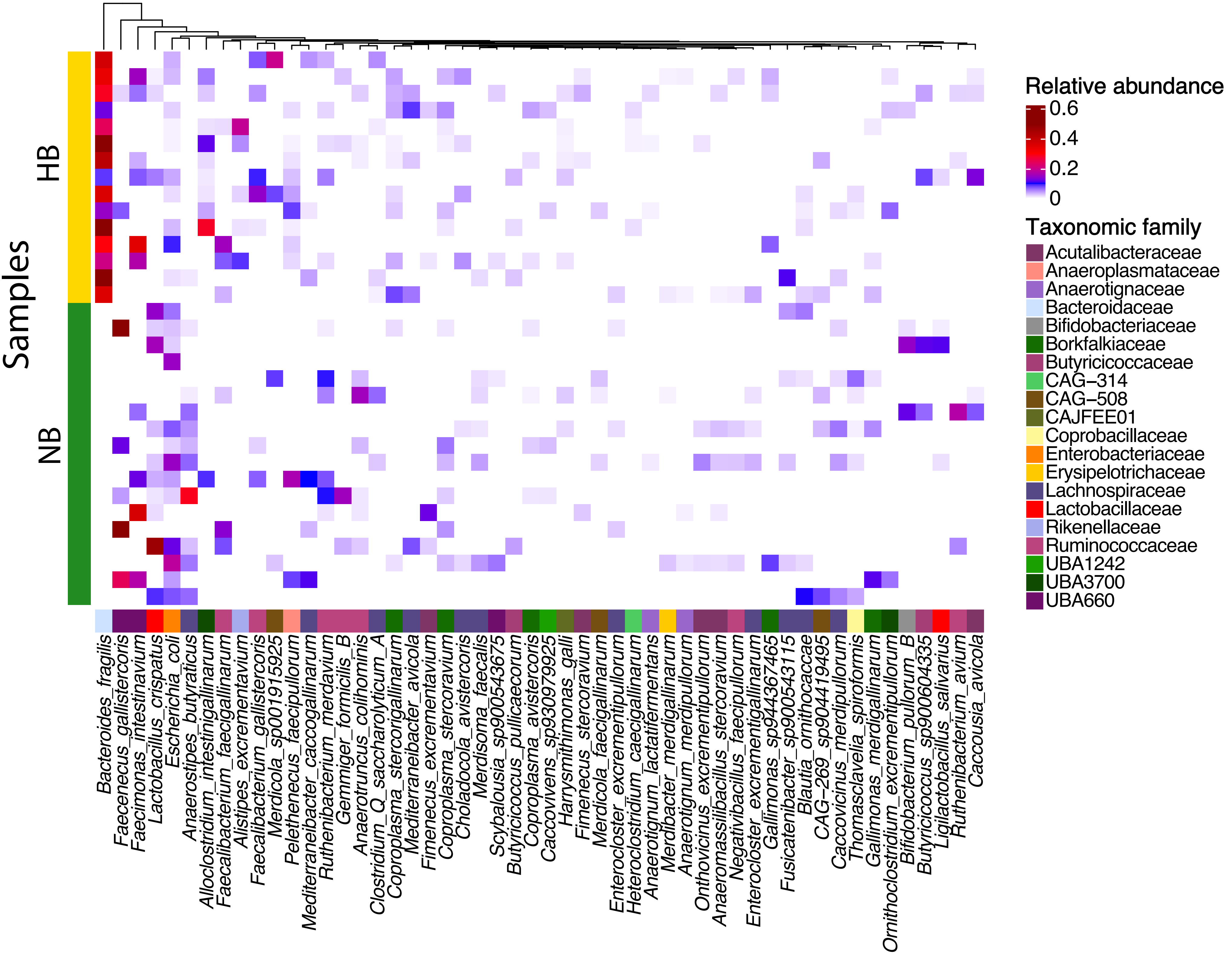

### Supplemental Figure 2

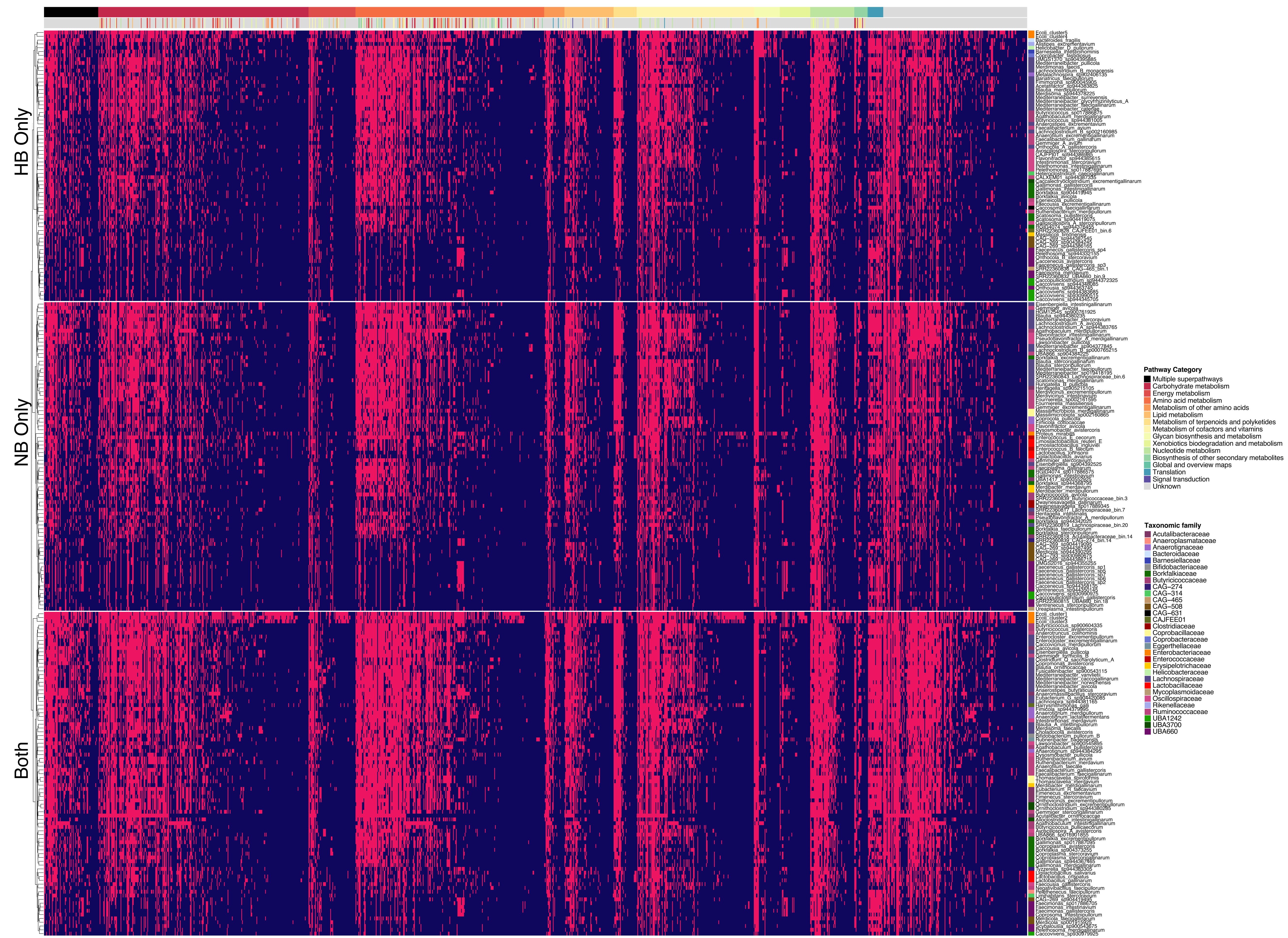

### Supplemental Figure 3

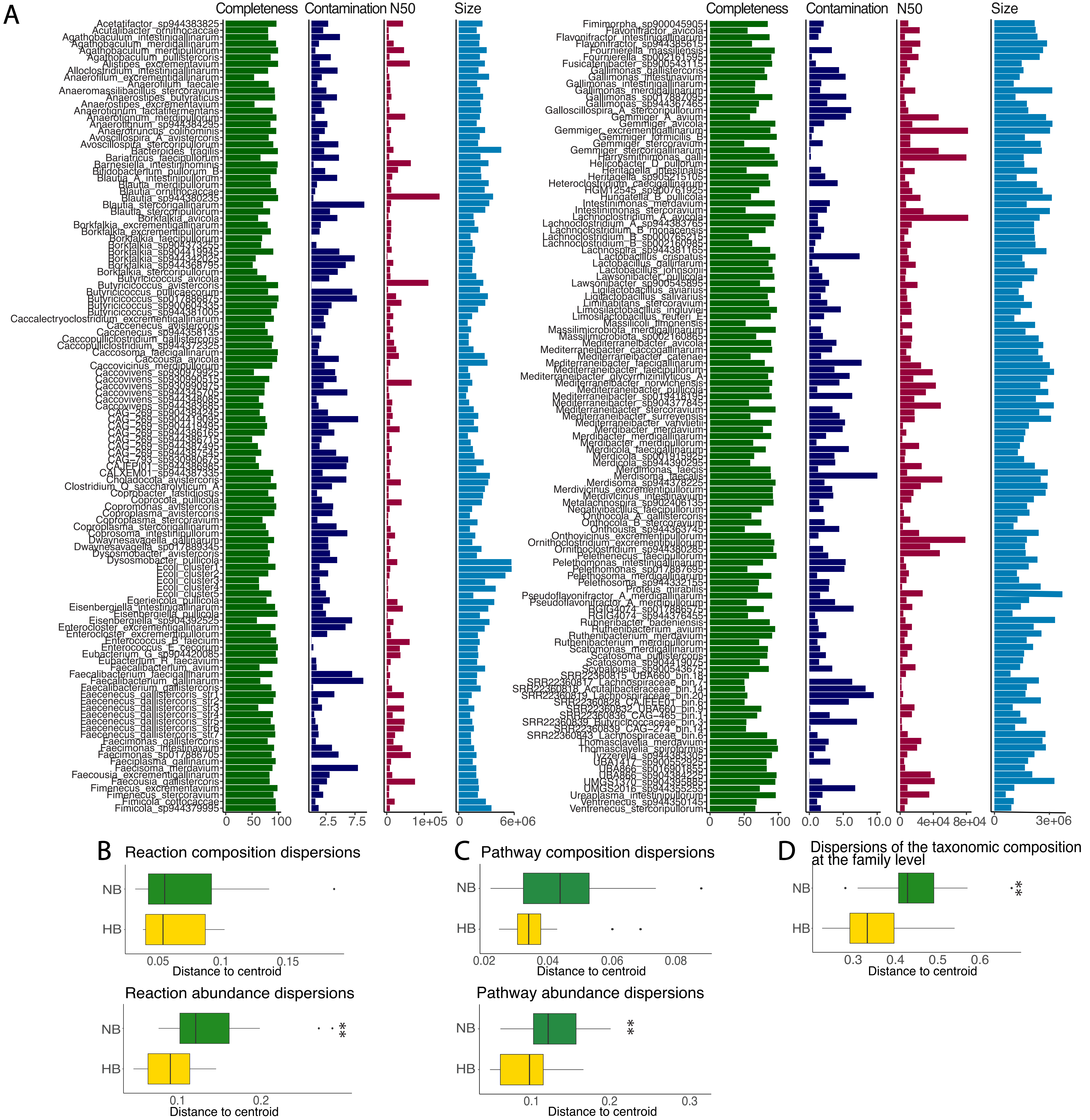

### Supplemental Figure 4

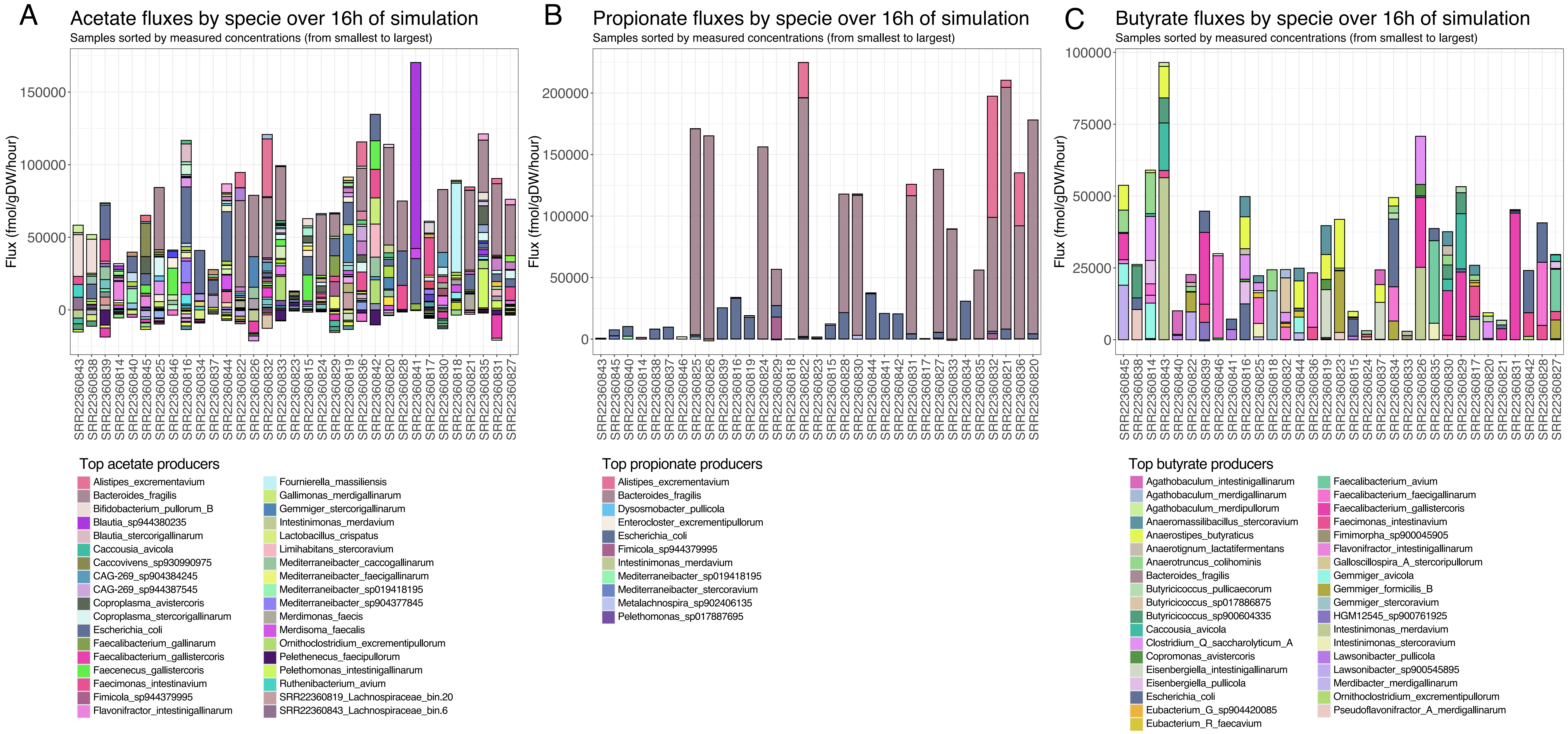

### Supplemental Figure 5

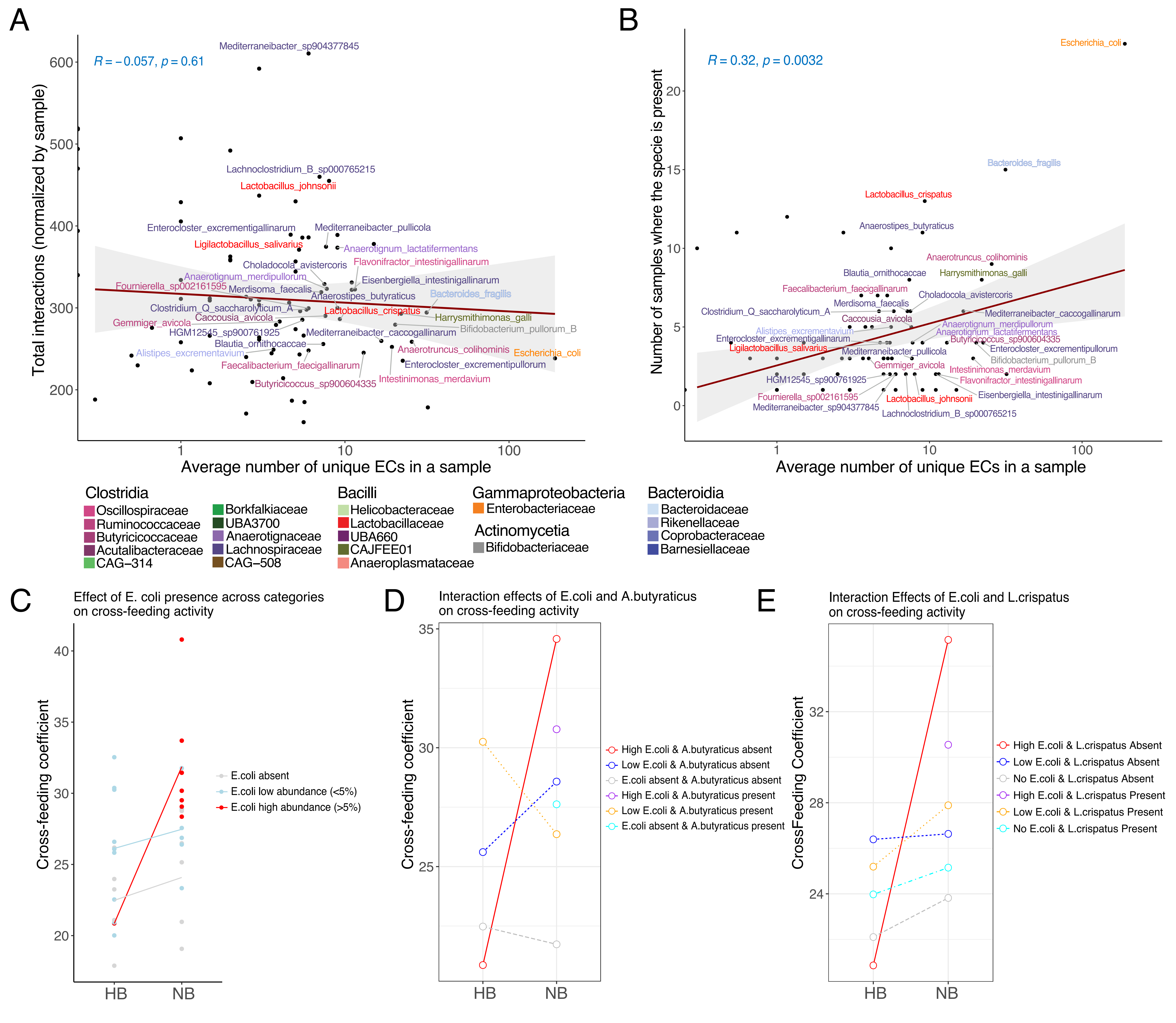

### Supplemental Figure 6

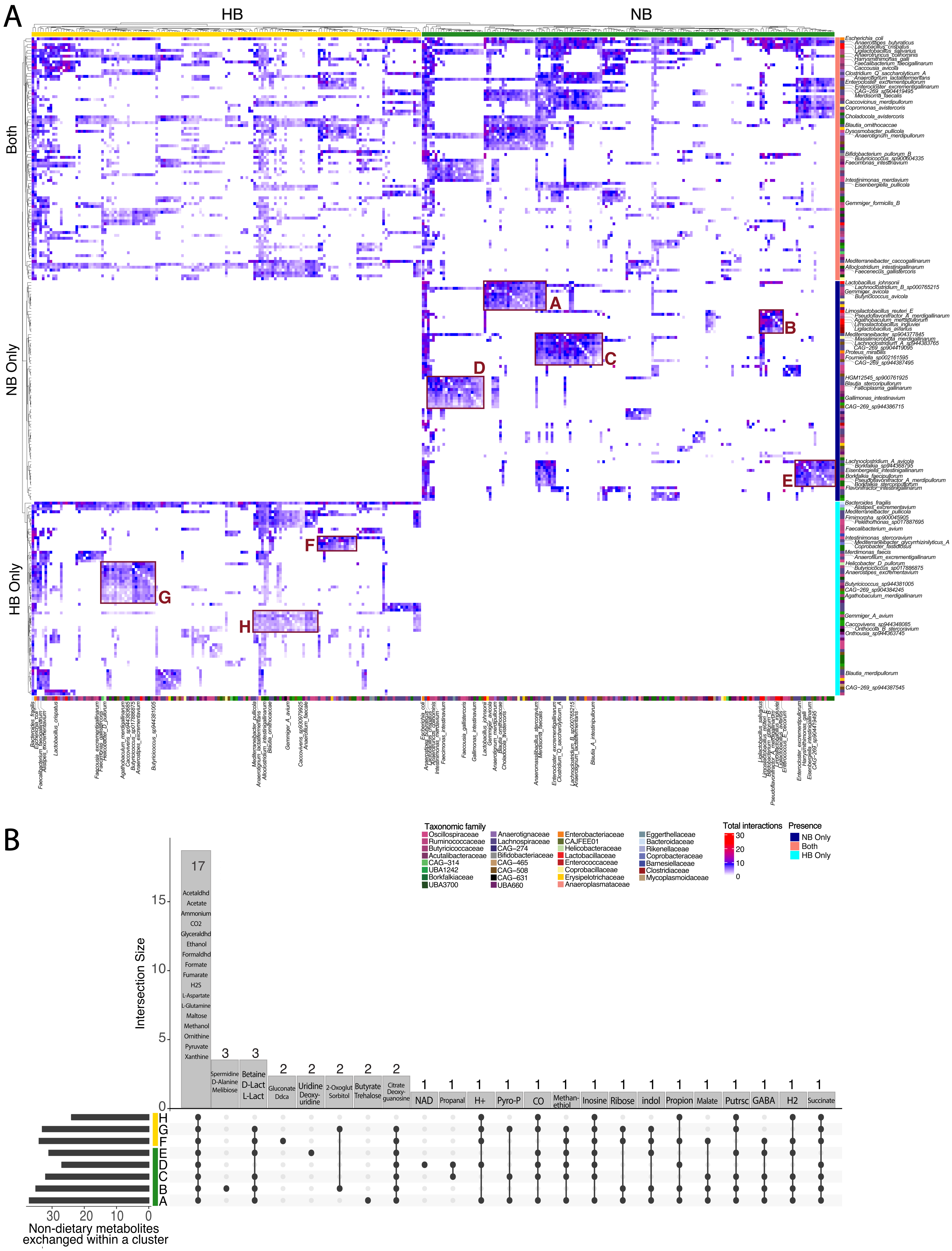
